# TDP-43 pathology triggers SRRM4-dependent cryptic splicing of G3BP1 in ALS/FTD

**DOI:** 10.1101/2025.09.17.676845

**Authors:** Hana Fakim, Asmita Ghosh, Emmanuel Amzallag, Yehuda M. Danino, Valérie Triassi, Anna-Leigh Brown, Nanu Pal, Jade-Emmanuelle Deshaies, Alicia Dubinski, Andréanne Lacombe, Carolane Fauchon, Arash Meghdadi Esfahani, Karen Ling, Frank Rigo, Paymaan Jafar-Nejad, NYGC ALS Consortium, Nicole J. Francis, Jean-François Trempe, Pietro Fratta, Alyssa N. Coyne, Eran Hornstein, Christine Vande Velde

## Abstract

Loss of nuclear TDP-43 is a defining feature of the neurodegenerative diseases amyotrophic lateral sclerosis (ALS) and frontotemporal dementia (FTD), yet how this leads to selective neuronal vulnerability is poorly understood. Here, using human iPSC-derived neurons and a large multi-omics dataset of ALS/FTD patients, we demonstrate that TDP-43 pathology induces the inclusion of an in-frame cryptic exon in human *G3BP1*. The resulting CRYPTIC G3BP1 protein contains an additional 10–amino acids within the highly conserved NTF2L domain, which acts as a dominant negative and disrupts stress granule dynamics. We further show that cryptic exon inclusion in *G3BP1* upon TDP-43 loss is enriched in neurons. Mechanistically, the loss of TDP-43 unmasks a binding site for the neuron-specific splicing regulator SRRM4 within intron 2 of G3BP1, enabling the inclusion of the cryptic exon. Collectively, our findings reveal that neuron-specific regulatory mechanisms intersect with TDP-43 -mediated splicing and suggest a mechanistic basis for the increased neuronal vulnerability observed in ALS/FTD.

## INTRODUCTION

Amyotrophic lateral sclerosis (ALS) is a fatal, late-onset neurodegenerative disease characterized by the loss of motor neurons, with no cure or effective therapy [1, 2]. Its molecular basis remains poorly understood due to clinical and genetic heterogeneity [3, 4]. Familial ALS (fALS, ∼10% of cases) is most often caused by autosomal dominant mutations in *C9ORF72*, *SOD1*, *TARDBP*, and *FUS*, while the remaining 90% are sporadic (sALS) with largely unknown genetic causes [3–6]. Importantly, 97% of all ALS patients exhibit nuclear-to-cytoplasmic mislocalization and aggregation of TAR DNA-binding protein 43 (TDP-43), despite mutations in its encoding gene accounting for only 2–5% of fALS [3–7]. This TDP-43 pathology also occurs in ∼45% of frontotemporal dementia (FTD) cases and other neurodegenerative disorders, including Alzheimer’s disease, Limbic-predominant age-related TDP-43 encephalopathy (LATE), and inclusion body myopathy, collectively termed ‘TDP-43 proteinopathies’ [6, 8].

TDP-43 is a ubiquitously expressed and predominantly nuclear RNA/DNA binding protein with multiple functions in RNA processing such as mRNA splicing, stability, transport, translation and miRNA biogenesis [8, 9]. TDP-43 has emerged as a key splicing regulator, where it binds uracil-guanine (UG) rich motifs and functions as a splicing repressor. TDP-43 loss-of-function *in vitro* and in patients results in the de-repression of aberrant splice sites and inclusion of intronic sequences, referred to as ‘cryptic exons’, into mature transcripts [10, 11]. Importantly, TDP-43 redistribution within the nucleus, which precedes cytoplasmic mislocalization, has been linked with cryptic splicing and is detectable prior to clinical symptom onset in ALS patients [12]. This suggests that early TDP-43 loss-of-function events, such as cryptic splicing, may play a pivotal role in driving ALS pathogenesis.

The inclusion of cryptic exons can lead to premature termination codons that trigger nonsense-mediated decay (NMD) or premature polyadenylation sites, as described in *UNC13A* [13, 14] and *STMN2* [15–17], respectively. Cryptic exon inclusion in *STMN2* results in a reduction in its transcript and protein levels, leading to axonal regeneration deficits [15, 17]. Less frequently, the inclusion of cryptic exons occurs in-frame and translates novel protein isoforms, such as in *HDGFL2* [18] or *MYO18A* [19]. How these cryptic protein isoforms influence the normal function of their respective proteins is not known. Intriguingly, cryptic splicing arising from TDP-43 loss-of-function varies considerably across cell types [10, 20], yet the molecular basis for this is not understood. RNA splicing is a highly complex process regulated by enhancer and repressor *cis*-acting elements that modulate spliceosome recruitment and splice site selection [21]. Many of the splicing factors recruited to these elements exhibit tissue/cell type-specific expression patterns [23, 24]. How these splicing factors compete/co-operate in TDP-43-mediated splicing repression remains to be elucidated and will be key to understanding the complexity and consequences of cryptic splicing.

Given ALS’s late-onset and sporadic nature, a combination of environmental stresses and genetic predisposition are posited to drive pathogenesis [22]. Accumulating evidence suggests that stress granules, cytoplasmic condensates composed of non-translating messenger ribonucleoproteins (mRNPs) that assemble as a survival response to stress [23], may play a key role in ALS. Specifically, some studies report that stress granules prevent TDP-43 aggregation and mitigate neurodegeneration [24–26], whereas others propose that they drive TDP-43 aggregation [27, 28].

We have previously shown that TDP-43 binds at the 3’ UTR to stabilize the mRNA encoding for G3BP1 (Ras-GTPase-activating protein-binding protein 1) [29, 30]. G3BP1 is a highly conserved RNA-binding protein (RBP) best known for its critical role in stress granule assembly under multiple cellular stresses. Further investigation is required to clarify the role of stress granules in ALS and across different stages of the disease.

In this study, we demonstrate that TDP-43 depletion in human iPSC-derived cortical neurons as well as TDP-43 pathology in ALS/FTD patient-derived motor neurons and post-mortem brains results in the inclusion of a cryptic exon in human *G3BP1*. This cryptic exon in *G3BP1* occurs in-frame within the NTF2L dimerization and protein-interaction domain of G3BP1. We report a reduction in the ability of CRYPTIC G3BP1 to interact with stress granule proteins compared to wild type (WT). Consistent with these observations, CRYPTIC G3BP1 abolishes stress granule formation during oxidative stress and functions as a dominant negative. We further establish that cryptic exon inclusion in *G3BP1* upon TDP-43 loss is enriched in neurons and mechanistically attribute this to the competition between TDP-43 and the neuronal splicing enhancer SRRM4 (splicing regulator serine/arginine repetitive matrix 4). Thus, TDP-43 loss-of-function permits SRRM4-dependent spurious splicing, producing an alternatively spliced CRYPTIC G3BP1 that impairs protective stress granule assembly and potentially contributes to neuronal vulnerability observed in TDP-43 proteinopathies. Collectively, this work sheds lights on the complexity of cryptic splicing in ALS/FTD and implicates G3BP1 as a promising therapeutic target.

## RESULTS

### 1. TDP-43 depletion in cortical neurons leads to the inclusion of a cryptic exon in *G3BP1*

We have previously reported that TDP-43 stabilizes *G3BP1* mRNA by binding to its 3′ UTR [30]. In searching TDP-43 eCLIP (enhanced nucleotide resolution UV crosslinking and immunoprecipitation) datasets [31], we identified additional TDP-43 binding sites located within intron 2 of human *G3BP1* (**Figure S1a**). Given the established role for TDP-43 in mRNA splicing [10, 32], we examined whether TDP-43 regulates the pre-mRNA splicing of *G3BP1*. We analyzed differential splicing in a publicly available RNA-seq dataset of human induced pluripotent stem cell (iPSC)-derived cortical neurons (i^3^-cortical neurons) depleted for TDP-43 using CRISPR inhibition versus control [13]. Intriguingly, the loss of TDP-43 resulted in a 30 nucleotide in-frame cryptic exon inclusion between exons 2 and 3 of *G3BP1* (hg38: chr5:151787765-151787794), that we termed exon2a (**Figure 1a-b, S1b**). The surrounding sequence shows that *G3BP1* exon2a is flanked by canonical AG/GT 3′/5′ splice sites and features two consensus TDP-43 binding sites downstream of the 5′ splice site that align with TDP-43 eCLIP binding peaks, supporting a direct regulation by TDP-43 (**Figure 1b**, **S1a**). Our sequence alignment analysis of *G3BP1* exon2a demonstrates that it is highly conserved across numerous primates but is not present in mice nor other vertebrates (**Figure S2 a-b**). *G3BP1* exon2a also shows a lack of sequence conservation and the absence of cryptic 3′/5′ splice sites in its paralog *G3BP2* (**Figure S2c**).

**Figure 1.**
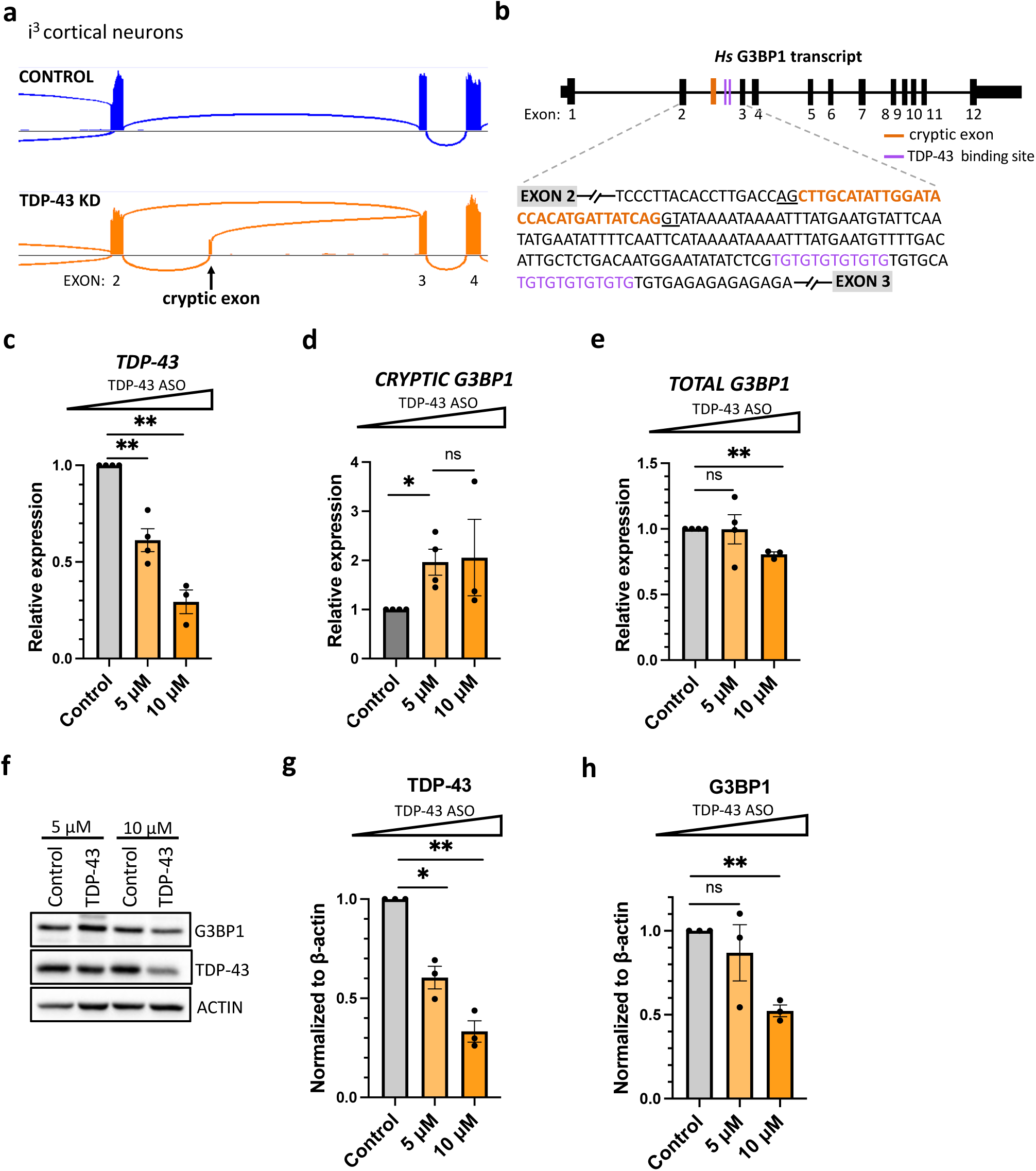
A cryptic exon inclusion event occurs in *G3BP1* transcript upon TDP-43 KD in i^3^cortical neurons. (a) Representative sashimi plot showing cryptic exon inclusion event (arrow) between exon 2 and 3 of human *G3BP1* transcript in TDP-43 depleted iPSC-derived cortical i^3^Neurons. (b) Schematic of cryptic exon location in human *G3BP1*. 30 nucleotide cryptic exon (exon2a) sequence is shown in orange. Two downstream TDP-43 consensus binding sites from eCLIP dataset shown in purple. (c-e) RT-qPCR analysis for specified gene (normalized to housekeeping genes *GAPDH* and *18S*) as a fold change over control. x axis shows μmolar concentration of TDP-43 ASO or control (10μM non-targeting ASO) treatment condition. Data are mean ± s.e.m; *n* = 3-4 biological replicates; statistical analysis performed is multiple unpaired t test with Welch’s correction. ns =not significant, ^∗^ < 0.05, ^∗∗^ < 0.01. (f) Representative western blot showing levels of G3BP1 and TDP-43 in i^3^cortical neurons treated with 5 μM or 10 μM of TDP-43 ASO versus 10 μM non-targeting control ASO. Actin was used as a loading control. (g-h) Protein levels quantified by western blot (normalized to actin loading control) as a fold change to control non-targeting ASO. x axis shows μmolar concentration of TDP-43 ASO or control (10μM non-targeting ASO) treatment condition. Data are mean ± s.e.m.; *n* = 3 biological replicates; statistical analysis performed is multiple unpaired t test with Welch’s correction. ns = not significant, ^∗^ < 0.05, ^∗∗^ < 0.01.

To further assess the link between TDP-43 depletion and exon2a inclusion in *G3BP1,* we performed a dose-dependent depletion of TDP-43 using 5 μM and 10 μM of RNase H-dependent antisense oligonucleotides (ASOs) targeting TDP-43 in i^3^-cortical neurons. ASO treatments resulted in ∼40% and ∼70% knockdown of *TDP-43* compared to 10 μM non-targeting ASO treatment, respectively, as assessed by RT–qPCR (**Figure 1c**). Using primers designed against the *G3BP1* exon2-2a boundary (**Figure S2d**), we observed a ∼2-fold increase in the inclusion of the cryptic exon in *G3BP1* in neurons treated with 5 μM TDP-43 ASO, with no change in total *G3BP1* levels (**Figure 1d, e**). At 10 μM ASO treatment, we noted an increase in *CRYPTIC G3BP1* similar to the 5 μM treatment, yet in addition observed a ∼20% decrease in total *G3BP1* mRNA (**Figure 1d, e**) and ∼40% reduction in protein levels (**Figure 1f-h**). Exon2a inclusion in *G3BP1* occurs in-frame and therefore does not trigger non-sense-mediated (NMD) decay, thus the latter is consistent with a disruption in TDP-43-mediated stabilization at the 3′ UTR, as we reported previously [30]. Taken together, our findings suggest that TDP-43 regulates human *G3BP1* mRNA through two mechanisms: by repressing inclusion of the cryptic exon2a and by stabilizing the *G3BP1* mRNA via the 3′ UTR regulatory element.

### 2. *CRYPTIC G3BP1* is detected in ALS/FTD patient neurons with TDP-43 pathology

To assess exon2a inclusion in *G3BP1* in the context of disease, we utilized iPSC lines from the Answer ALS (AALS) consortium that were derived from ALS patients and non-neurological controls [33]. These iPSC lines were differentiated into spinal motor neurons (iMNs) [33]. sALS lines were stratified by TDP-43 pathology using a previously characterized TDP-43 loss-of-function splicing signature based on 14 gene expression/splicing alterations [34]. We observed that the expression of *CRYPTIC G3BP1* was increased ∼4-fold in sALS iMNs with TDP-43 pathology relative to sALS iMNs without TDP-43 pathology or control lines, as assessed by RT-qPCR (**Figure 2a**). Elevated expression of *CRYPTIC G3BP1* in iMNs with TDP-43 mutations was increased ∼6-fold compared to control, however, likely due to the small sample size (n=3) we did not reach statistical significance (**Figure 2a**). *CRYPTIC G3BP1* levels were ∼5-fold higher in iMNs harboring *C9ORF72* repeat expansion compared to control, but no change was observed in iMNs with *SOD1* mutations (**Figure 2a**), consistent with TDP-43 being mislocalized in *C9ORF72-ALS,* but not in *SOD1-ALS* [35]. These results suggest that the cryptic splicing of *G3BP1* is confined to ALS clinical subtypes where TDP-43 pathology is reported to occur.

**Figure 2:**
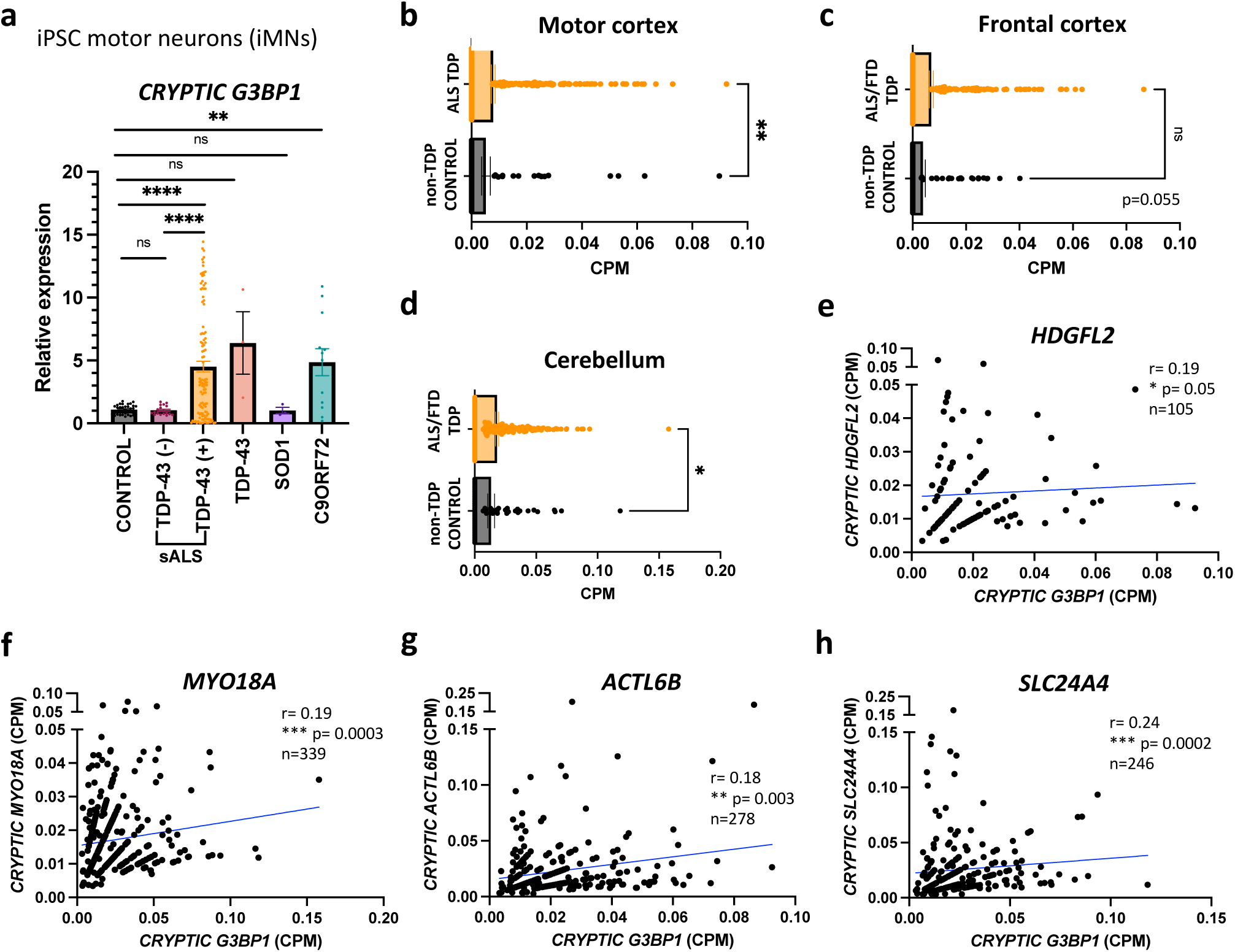
*CRYPTIC G3BP1* is detected in ALS/FTD patient neurons with TDP-43 pathology. (a) RT-qPCR analysis for *CRYPTIC G3BP1* mRNA normalized to housekeeping genes *GAPDH* and *18S* in Answer ALS patient-derived iPSC motor neurons (iMNs). Sporadic ALS (sALS) patient lines were sorted into TDP-43 (-) versus TDP-43 (+) based on TDP-43 splicing signature. Control n= 33, sALS TDP-43 (-) n= 18, sALS TDP-43 (+) n= 111, TDP-43 n= 3, SOD1 n=3, C9ORF72 n= 12. Data are mean ± s.e.m; statistical analysis performed is multiple unpaired t test with Welch’s correction. ns = not significant, ^∗^ < 0.05, ^∗∗^ < 0.01, ^∗∗∗^ < 0.001, ^∗∗∗∗^ < 0.0001. (b-d) Expression of *CRYPTIC G3BP1* transcript across indicated brain regions from NYGC ALS Consortium RNA seq depicted as counts per million (CPM). Samples were stratified by TDP-43 pathology, comparing ALS/FTD patients with TDP-43 pathology (ALS/FTD TDP) to TDP-43 pathology-free samples (non-neurological controls or patients with *SOD1* or *FUS* mutations (non-TDP CONTROL)). Motor cortex (non-TDP CONTROL n=89, ALS TDP n=445), Frontal cortex (non-TDP CONTROL n=106, ALS/FTD TDP n=343), Cerebellum (non-TDP CONTROL n=66, ALS/FTD TDP n=274). Data are mean ± s.e.m; statistical analysis performed is Mann-Whitney test. ns = not significant, ^∗^ < 0.05, ^∗∗^ < 0.01. (e-h) Correlation between *CRYPTIC G3BP1* levels and indicated cryptic exon-containing transcripts (in CPM) within the central nervous system (CNS) of ALS/FTD patients with TDP-43 pathology. Statistical analysis performed is Spearman correlation test. ns =not significant, ^∗^ < 0.05, ^∗∗^ < 0.01, ^∗∗∗^ < 0.001.

To detect *G3BP1* exon2a inclusion in ALS/FTD patient brains, we examined RNA-seq data from the New York Genome Center (NYGC) ALS Consortium encompassing transcriptomics data from 2361 tissues from 640 individuals with ALS or FTD versus non-neurological controls. To evaluate the impact of TDP-43 pathology on cryptic splicing of *G3BP1*, we performed an analysis comparing patients with TDP-43 pathology to pathology-free samples, that were either of controls or of patients harboring *SOD1* or *FUS* mutations (non-TDP controls). Consistent with known TDP-43 mislocalization in these regions [36], we observed higher levels of *CRYPTIC G3BP1* in the motor cortex and an increasing trend (p=0.055) in the frontal cortex of ALS/FTD patients with TDP-pathology compared to non-TDP controls (**Figure 2b, c**). No difference in *CRYPTIC G3BP1* levels was observed in the hippocampus, temporal cortex, or spinal cord (**Figure S3a-c**). The absence of *CRYPTIC G3BP1* in the spinal cord is unexpected and may reflect the severity of TDP-43 pathology and neuronal loss in this region [37]. We also noted an increased expression of *CRYPTIC G3BP1* in the cerebellum of ALS/FTD patients with TDP-pathology compared to non-TDP controls (**Figure 2d**), a region more recently implicated in ALS/FTD [38, 39]. Finally, *CRYPTIC G3BP1* levels positively correlated with several in-frame TDP-43-induced cryptic exons such as in *HDGFL2*, *MYO18A* and *ACTL6B* as well as frameshift cryptic exon in *SLC24A4* (**Figure 2e-h**) and *RAP1GAP* (**Figure S3d**). Interestingly, *CRYPTIC G3BP1* did not correlate with cryptic exons in *STMN2*, *UNC13A* and *PFKP* (**Figure S3e-g**), suggesting possible additional splicing regulation or sensitivity to TDP-43 pathology.

Collectively, these data reveal a strong relationship between TDP-43 pathology and *CRYPTIC G3BP1*, both in cell culture and in ALS/FTD neuropathology, supporting a direct role for TDP-43 in repressing cryptic exon2a in *G3BP1*.

### 3. *CRYPTIC G3BP1* adds 10 amino acids in-frame in NTF2L domain and alters its interactome

The inclusion of cryptic exon2a, being a multiple-of-three insertion, is predicted to maintain the G3BP1 reading frame. G3BP1 is composed of 466 amino acids (aa) and contains an N-terminal NTF2L (nuclear transport factor 2-like) domain, two central intrinsically disordered regions (IDRs) comprised of acidic and proline-rich (PxxP) domains, and the C-terminal RNA-binding (RRM) and arginine-glycine rich (RGG) motifs (**Figure 3a**). The CRYPTIC peptide encoded by exon2a inserts 10 aa within the highly conserved NTF2L domain of G3BP1 (**Figure 3a**). The G3BP1 NTF2L domain enables its dimerization [40] and is required for stress granule assembly by mediating pivotal protein-protein interactions that promote oligomerization [41, 42]. The cryptic peptide inserts after histidine 31 (H31) in G3BP1 adjacent to phenylalanine 33 (F33), one of three phenylalanine residues that form a hydrophobic pocket mediating interactions with short linear motifs (SLiMs) in the key stress granule proteins CAPRIN1, UBAP2L and USP10 [43–45].

**Figure 3:**
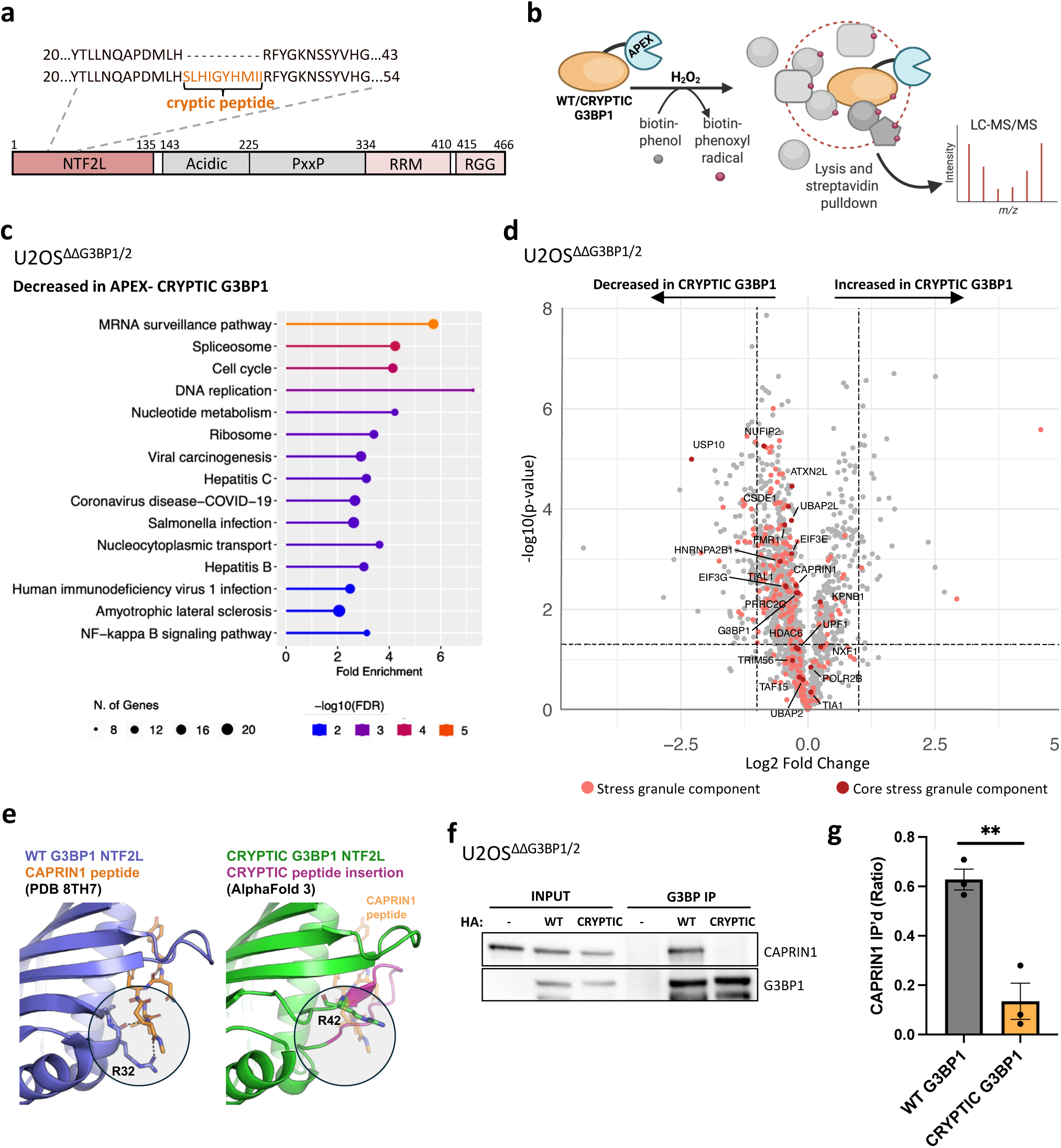
The cryptic exon in *G3BP1* leads to the addition of 10 amino acids in-frame within the NTF2L domain. (a) Schematic of domain architecture of human G3BP1 protein, encompassing NTF2L domain, acidic motif, proline-rich (PxxP) motif, RNA recognition motif (RRM) and arginine-glycine rich (RGG) motif. CRYPTIC peptide inserts into NTF2L domain after residue H31. (b) Schematic representation of APEX proteomics assay. APEX-tagged WT or CRYPTIC G3BP1 were stably expressed in U2OS^ΔΔG3BP1/2^ cells. The addition of H_2_O_2_ induces APEX peroxidase activity, resulting in biotin-phenoxyl (BP) radicals that tag proximal proteins. Cells are then lysed under harsh conditions and biotinylated proteins are purified using streptavidin resin and analyzed by mass spectrometry. (c) KEGG pathway analysis of significantly downregulated genes (p< 0.05) in CRYPTIC G3BP1-APEX versus WT G3BP1-APEX samples, after normalizing to NES-APEX, using ShinyGO tool. (d) Volcano plot from Figure S5a highlighting all stress granule protein components in light red. “Core” stress granule proteins are labelled and depicted in dark red. (e) Comparison of the crystal structure of NTF2L domain of WT G3BP1 bound to CAPRIN1 (violet/orange, left) versus CRYPTIC G3BP1 (green, right) predicted with Alphafold 3. Both structures were superposed and are shown in the same orientation. The cryptic peptide insertion is shown in magenta, and the CAPRIN1 peptide is superposed as transparent sticks on the CRYPTIC G3BP1 structure. (f) Western blot analysis probed for indicated proteins showing input and co-immunoprecipitated proteins using G3BP1 antibody in lysates from U2OS^ΔΔG3BP1/2^ cells transiently transfected with HA-WT or CRYPTIC G3BP1. Untransfected cells were used as a negative control. (g) Quantification of CAPRIN1 intensity normalized to G3BP1 in CO-IP. *n* = 3 biological replicates; statistical analysis performed is multiple unpaired t test with Welch’s correction. ns = not significant, ^∗^ < 0.05, ^∗∗^ < 0.01.

To assess the impact of the CRYPTIC peptide on the G3BP1 interactome under basal stress-free conditions we employed APEX2-proximity labelling and proteomics [46, 47] (**Figure 3b**). To avoid confounding effects from endogenous G3BP1/G3BP2 and TDP-43–mediated regulation of *G3BP1* at its 3′ UTR, we expressed APEX2-tagged WT [47] or CRYPTIC G3BP1 in a U2OS cell line CRISPR-edited to lack G3BP1 and G3BP2 (U2OS^ΔΔG3BP1/2^). APEX-G3BP1 constructs were stably expressed under doxycycline-inducible control at comparable levels (**Figure S4a**) and APEX2 fused to a nuclear export signal (NES) was used a cytosolic bait control. Cells were treated with a short pulse of hydrogen peroxide (H₂O₂) that radicalizes pre-incubated biotin-phenol (BP) to covalently tag proteins within the vicinity of APEX2-G3BP1 or APEX2-NES (**Figure 3b**). Biotinylated proteins were then pulled down using streptavidin beads and the unbiased proteome was analysed by mass spectrometry (as in [47]). We confirmed that endogenous proteins were tagged with BP upon APEX2 activation by western blot (**Figure S4b**) and noted a strong correlation across experimental replicates (**Figure S4c**). No-BP lysates were used as negative controls to exclude proteins non-specifically bound to streptavidin beads. Candidate G3BP1-interacting proteins were identified by applying Student’s t tests (false discovery rate [FDR] cutoff p< 0.05) to proteins associated with APEX-G3BP1 samples, relative to APEX-NES samples. Principle component analysis (PCA) revealed distinct proteomic signatures between APEX-WT G3BP1 and APEX-CRYPTIC G3BP1 (**Figure S4d**). A summary of the proteomic dataset generated is provided in **Table S1.**

When compared to APEX-WT G3BP1, we identified 688 proteins that were significantly reduced (p value <0.05) (102 with a >2-fold enrichment) and 261 proteins that were enriched (37 with >2-fold enrichment) in APEX-CRYPTIC G3BP1 (**Figure S5a**). Gene Ontology (GO) enrichment analysis revealed that the protein interactions reduced in CRYPTIC G3BP1 compared to WT were enriched for RNA metabolism pathways, including mRNA surveillance, spliceosome, and ribosome-related processes (**Figure 3c**). Conversely, although less abundant, protein interactions that increased with CRYPTIC G3BP1 compared to WT were associated with ER processing and neurogenerative diseases (**Figure S5b**). These changes are unlikely due to the mislocalization of CRYPTIC G3BP1, as expression of HA- and YFP-tagged CRYPTIC G3BP1 localization was comparable to WT G3BP1 expressed in U2OS^ΔΔG3BP1/2^ and i^3^-cortical neurons, respectively (**Figure S5c**). Together, these results suggest that the CRYPTIC peptide in G3BP1 rewires its protein interactome compared to WT, likely altering its cellular functions.

Stress granule assembly is driven by interactions within a core protein–RNA network, comprised of approximately 36 “core” proteins and their associated RNAs, which trigger liquid-liquid phase separation (LLPS) upon reaching a threshold concentration [41, 42]. G3BP1 and G3BP2 are central nodes within this network due to their high valency, the capacity to engage in numerous simultaneous interactions, and thus are essential for the formation of stress granules during oxidative stress [41, 42]. To assess the ability of CRYPTIC G3BP1 to interact with the stress granule network, we compared our APEX results with a published list of 489 proteins localized to stress granules in U2OS^ΔΔG3BP1/2^ under oxidative stress [47]. Of those that overlapped, most of these stress granule components were reduced in APEX-CRYPTIC G3BP1 compared to WT (142 proteins), while only 22 stress granule components displayed an increased interaction with CRYPTIC G3BP1 (**Figure 3d**). Among the 36 proteins that comprise the stress granule core complex [41], interactions with 12 (ATXN2L, CAPRIN1, CSDE1, EIF3E, EIF3G, FMR1, HNRNPA2B1, NUFIP2, PRRC2C, TIAL1, UBAP2L, and USP10) were reduced, whereas only one (KPNB1) exhibited increased interaction.

As a published crystal structure for CAPRIN1-G3BP1 NTF2L domain exists [48], and given the ascribed role for CAPRIN1 as an enhancer of stress granule formation [49], we chose to further explore the impact of the cryptic peptide on this interaction. Modelling of the CRYPTIC G3BP1 NTF2L structure using AlphaFold 3 [50] shows that the cryptic peptide is located in a short loop of the NTF2L domain and forms a longer loop with an extended β-strand conformation (**Figure 3e**). A superposition of the G3BP1-CAPRIN1 crystal structure (PDB 8TH7) on the CRYPTIC G3BP1 NTF2L structure predicted shows that the cryptic peptide in G3BP1 is positioned within the CAPRIN1 binding site, likely disrupting its interaction (**Figure 3e**). In particular, the crystal structure shows that both the main chain carbonyl and side chain of arginine 32 (R32) in G3BP1 form hydrogen bonds with CAPRIN1 that would be disrupted by the cryptic peptide located just before this residue (R42 in CRYPTIC G3BP1) (**Figure 3e**). We validated this experimentally by transiently transfecting U2OS^ΔΔG3BP1/2^ cells with HA-WT or CRYPTIC G3BP1 followed by co-immunoprecipitation using G3BP1 polyclonal antibody. Untransfected cells were used as a negative control. Herein, we observed a significant reduction in the ability of HA-CRYPTIC G3BP1 to co-immunoprecipitate CAPRIN1 compared to WT (**Figure 3f, g**).

Taken together, these results indicate that CRYPTIC G3BP1 exhibits a perturbed interactome compared to WT, disrupting its ability to interface with the stress granule protein network.

### 4. CRYPTIC G3BP1 alters stress granule formation

We tested whether CRYPTIC G3BP1 can form stress granules under external stress by transiently expressing GFP-tagged WT or CRYPTIC G3BP1 in U2OS^ΔΔG3BP1/2^ cells, which lack G3BP1/2 and therefore cannot form stress granules in response to oxidative stress (sodium arsenite) [41, 49]. Remarkably, U2OS^ΔΔG3BP1/2^ cells expressing GFP-tagged CRYPTIC G3BP1 remained devoid of stress granules in response to sodium arsenite (**Figure 4a, b**), whereas complementation of U2OS^ΔΔG3BP1/2^ cells with GFP-tagged WT G3BP1 rescued stress granule formation as previously reported [41, 49] (**Figure 4a, b**), suggesting that CRYPTIC G3BP1 is defective at nucleating stress granules. These results are consistent with CRYPTIC G3BP1’s reduced interaction with numerous core stress granule factors (**Figure 3d**), including CAPRIN1 (**Figure 3e-g**), suggesting a limited ability of CRYPTIC G3BP1 to attain the valency required for stress granule condensation [42].

**Figure 4:**
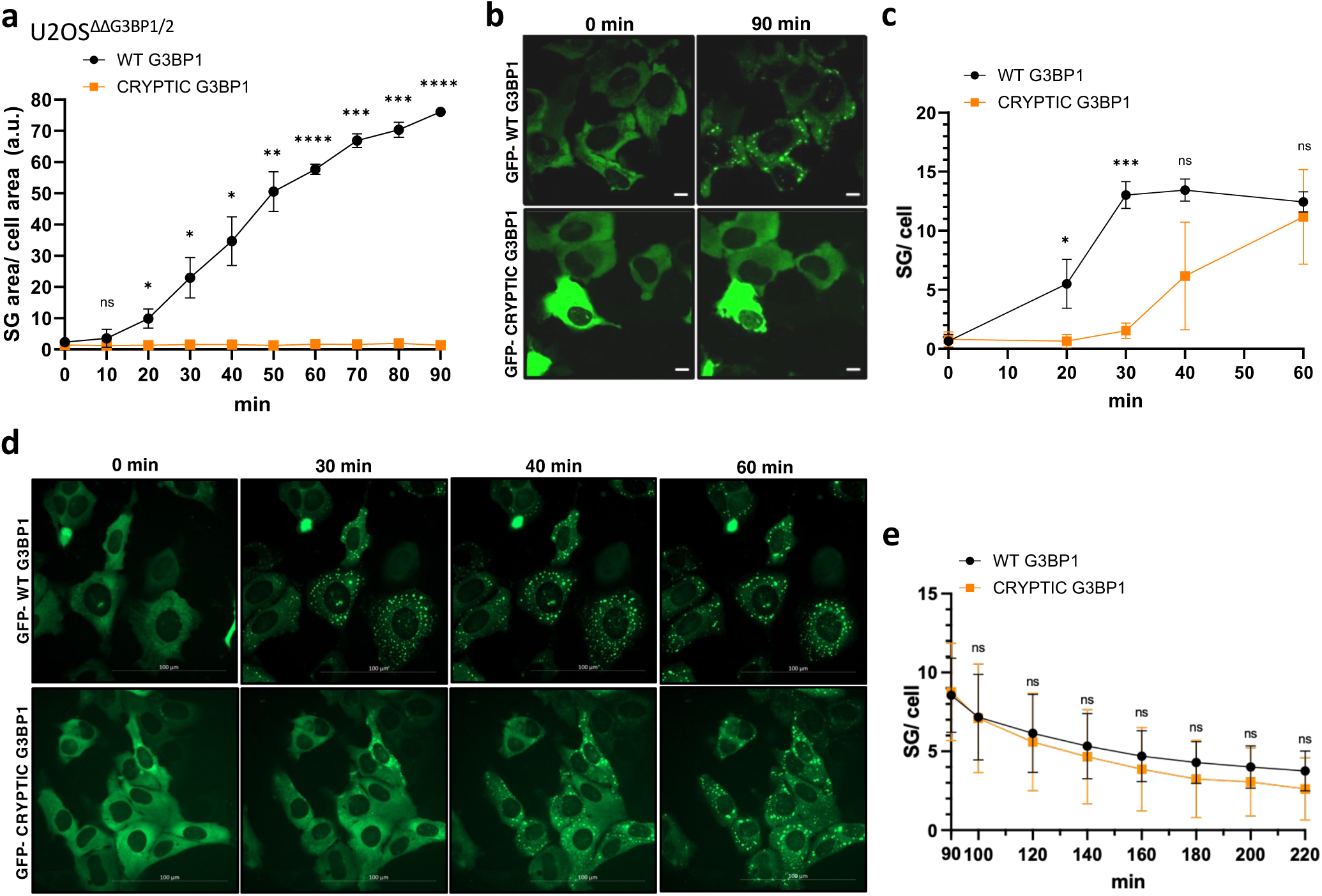
CRYPTIC G3BP1 alters stress granule formation. (a) Quantification of SG dynamics by live imaging of U2OS^ΔΔG3BP1/2^ cells transiently expressing GFP-WT G3BP1 or GFP-CRYPTIC G3BP1. Cells were subjected to 250 μM sodium arsenite for the indicated time. Datapoints depict SG area, normalized to cellular area (y axis), as a function of time (x axis). Data are mean ± SD; n = 3 biological replicates; statistical analysis performed is multiple unpaired t test with Welch’s correction. ∗ < 0.05, ∗∗ < 0.01, ∗∗∗ < 0.001, ∗∗∗∗ < 0.0001. (b) Representative images of live imaging results from (a) at indicated timepoints. Scale bar, 10 μm. (c) Quantification of SG dynamics by live imaging of U2OS^ΔΔG3BP1/2^ cells transiently expressing GFP-WT G3BP1 or GFP-CRYPTIC G3BP1. Cells were subjected to heat stress (43°C) for the indicated time. Datapoints depict SG number per cell, as a function of time (x axis). Data are mean ± SD; n = 3 biological replicates; statistical analysis performed is multiple unpaired t test with Welch’s correction. ∗ < 0.05, ∗∗ < 0.01, ∗∗∗ < 0.001, ∗∗∗∗ < 0.0001. (d) Representative images of live imaging results from (c) at indicated timepoints. Scale bar, 100 μm. (e) Quantification of SG disassembly dynamics by live imaging of U2OS^ΔΔG3BP1/2^ cells transiently expressing GFP-WT G3BP1 or GFP-CRYPTIC G3BP1. Cells were subjected to heat stress (43°C) for 90 mins followed by disassembly for the indicated time. Datapoints depict SG number per cell, as a function of time (x axis). Data are mean ± SD; n = 3 biological replicates; statistical analysis performed is multiple unpaired t test with Welch’s correction. ns=not significant.

Stress granule formation via heat stress does not require G3BP1 in U2OS^ΔΔG3BP1/2^ cells and therefore may serve as an orthogonal stress paradigm [41, 49, 51]. During heat stress (43°C), U2OS^ΔΔG3BP1/2^ complemented with GFP-tagged WT G3BP1 resulted in >10 stress granules per cell at 30 minutes, whereas this was significantly delayed until 60 minutes with CRYPTIC G3BP1 complementation (**Figure 4c, d**). Recovery dynamics from heat stress were not affected (**Figure 4e**). These results suggest that although G3BP1 is not required to nucleate stress granules during heat stress, the recruitment of CRYPTIC G3BP1 with reduced valency compared to WT alters stress granule condensation dynamics [42]. Taken together, these results demonstrate that CRYPTIC G3BP1 alters stress granule formation in response to both oxidative and heat stress.

### 5. CRYPTIC G3BP1 associates with WT G3BP1 and acts as a dominant negative

To interrogate the effect of CRYPTIC G3BP1 on the native protein, we investigated its ability to dimerize with WT G3BP1. The structure of the G3BP1 NTF2L domain has been previously solved using X ray crystallography [52], thus we used AlphaFold 3 [50] to predict the structure of the WT-CRYPTIC G3BP1 NTF2L dimer. These results show that the CRYPTIC peptide insertion occurs at opposite end of the G3BP1 dimerization interface (**Figure 5a and S5d**), likely maintaining interaction. To assess this, we transiently expressed GFP-tagged WT G3BP1 in combination with either HA-tagged WT or CRYPTIC G3BP1 in U2OS^ΔΔG3BP1/2^ cells and performed co-immunoprecipitation assays using anti-HA antibody. Expression of GFP-WT G3BP1 alone served as a negative control. Western blot analysis revealed that HA-CRYPTIC G3BP1 and HA-WT G3BP1 comparably co-immunoprecipitated with GFP-tagged WT G3BP1 (**Figure 5b**), suggesting that CRYTPIC G3BP1 is capable of associating with WT G3BP1.

**Figure 5.**
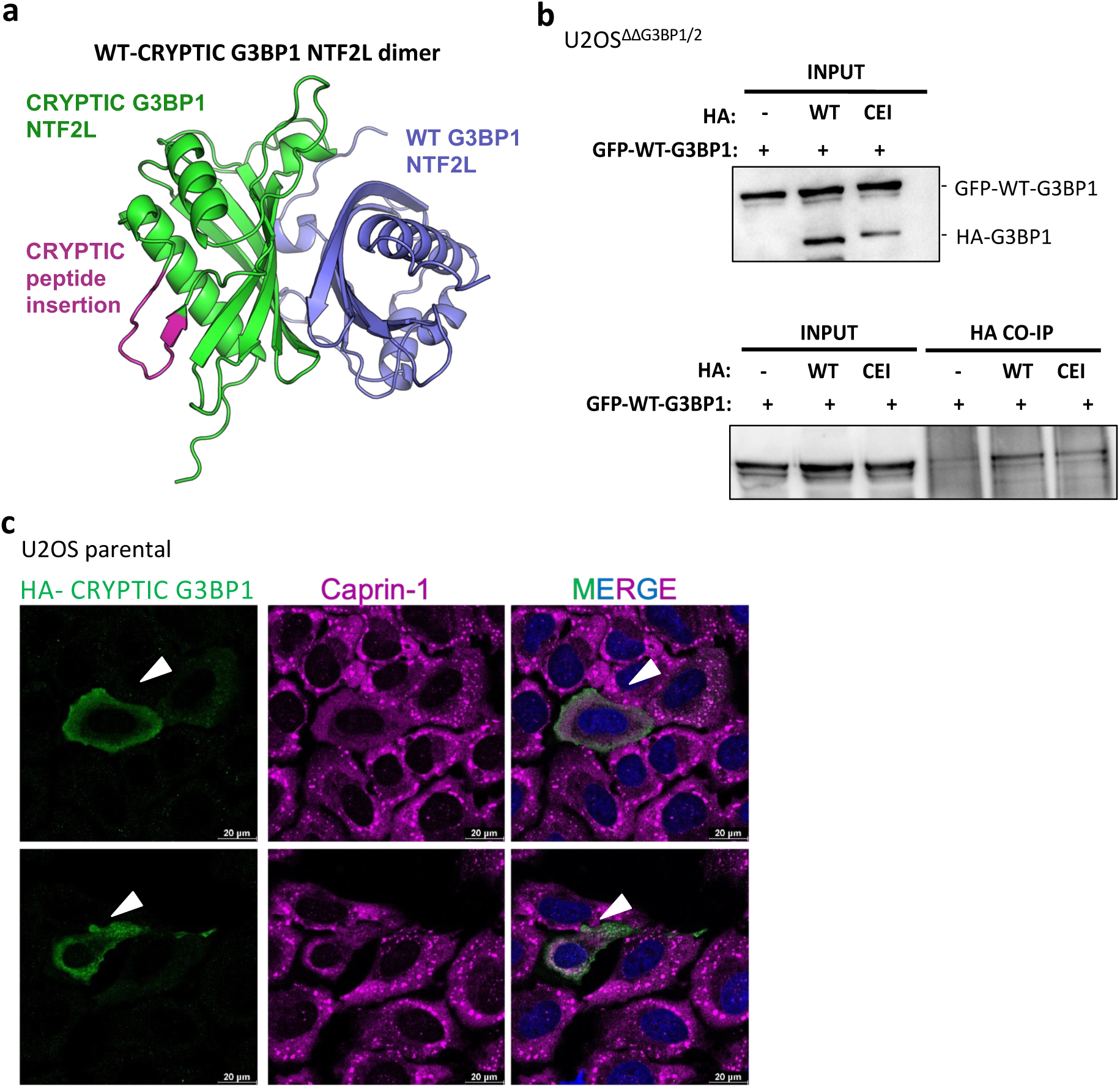
CRYPTIC G3BP1 associates with WT G3BP1 and acts as a dominant negative. (a) Structure of CRYPTIC G3BP1 (green) and WT G3BP1 (violet) NTF2L heterodimer predicted using Alphafold 3. CRYPTIC G3BP1 sequence peptide is depicted in magenta. (d) (Top) Western blot analysis probed for G3BP1 showing input levels of transiently transfected GFP-WT G3BP1 and HA-WT or CRYPTIC G3BP1 in U2OS^ΔΔG3BP1/2^ cells. (Bottom) Western blot analysis probed for GFP showing levels of GFP-WT G3BP1 co-immunoprecipitated using HA-antibody in cells expressing HA-WT G3BP1 versus HA-CRYPTIC G3BP1. Cells transfected with GFP-WT G3BP1 but no HA-tagged constructs was used negative control. (c) Immunofluorescence using CAPRIN1 antibody as a stress granule marker and HA to label HA-CRYPTIC G3BP1 transiently transfected (white arrowhead) U2OS parental cells. Cells were subjected to 500 μM sodium arsenite for 30 minutes.

Finally, to assess the impact of CRYPTIC G3BP1 on the WT protein’s function, we transfected low amounts of HA-tagged CRYPTIC G3BP1 into U2OS parental cells (containing endogenous G3BP1 and G3BP2). Cells transfected with CRYPTIC G3BP1 showed impaired stress granule formation upon oxidative stress relative to surrounding non-transfected cells (**Figure 5c**), supporting a dominant-negative function for CRYPTIC G3BP1. These results are consistent with the ability of CRYPTIC G3BP1 to interact with WT G3BP1, presumably reducing its ability to form large multimeric complexes with sufficient valency to nucleate stress granules.

### 6. Loss of TDP-43 unmasks an SRRM4 binding site required for inclusion of the cryptic exon in *G3BP1*

In non-neuronal cell lines such as U2OS and HeLa, we observed that depletion of TDP-43 using siRNA was insufficient to promote the inclusion of exon2a in *G3BP1*, despite cryptic exon inclusion in *UNC13A* (**Figure S6a-f**). To identify the missing splicing enhancer that drives cryptic splicing of *G3BP1* in iPSC-derived neurons and human brains in the context of TDP-43 pathology (**Figure 1, 2**), we searched for RBP-binding sites near the *G3BP1* cryptic exon. The neural-specific SR-related protein of 100 kDa (nSR100 or SRRM4, Ser/Arg-repeat matrix protein 4) emerged as a candidate RBP based on its restricted neuronal expression (GTEX [53], Protein Atlas [54], [55]) and the presence of multiple binding motifs (TGC) near the *G3BP1* cryptic splice sites and downstream TDP-43 binding region (**Figure S6g**). To assess the role of SRRM4 as the putative splicing enhancer, we generated a stable cell line expressing doxycycline-inducible FLAG-SRRM4 in U2OS cells, where endogenous SRRM4 expression is absent (**Figure 6a, S6h, Table S2**). Expression of SRRM4 promoted cryptic exon2a inclusion in *G3BP1* in a dose-dependent manner (**Figure 6b**), without changes in total *G3BP1* mRNA (**Figure S6i**). This effect was not due to changes in TDP-43 levels upon SRRM4 expression (**Figure 6c**). Conversely, we did not observe significant changes in endogenous *SRRM4* levels upon TDP-43 depletion in i^3^-cortical neurons (**Figure S6j**) and *SRRM4* remained undetected in U2OS or HeLa cells upon TDP-43 knockdown (**Table S2**). Interestingly, cryptic exon inclusion in *G3BP1* was increased up to ∼100-fold with 4 µg/ml dox and ∼5-fold in *STMN2* (**Figure 6b, d**), but had no effect on cryptic splicing of *UNC13A* (**Table S2**). These results are consistent with a role for SRRM4 in regulating the cryptic splicing of *G3BP1*, and to a lesser but significant degree enhancing cryptic exon inclusion in *STMN2*, in a neuron-specific manner.

**Figure 6:**
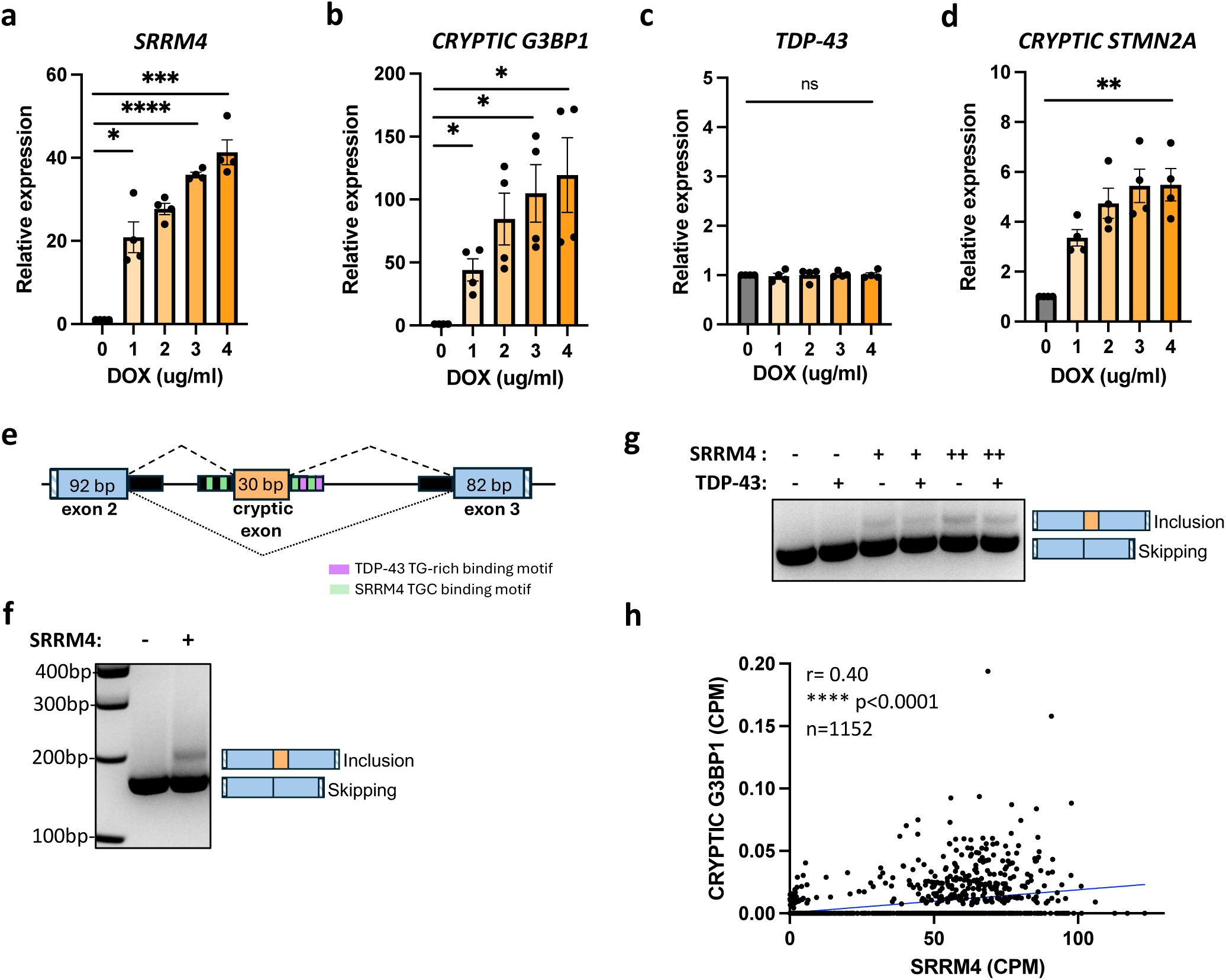
Loss of TDP-43 unmasks an SRRM4 binding site required for the inclusion of the cryptic exon in *G3BP1*. (a-d) RT-qPCR analysis for specified gene (normalized to housekeeping genes *GAPDH* and *18S*) as a fold change over no doxycycline (dox) control. x axis shows doxycycline concentration used to induce FLAG-SRRM4 expression in stable U2OS cell line. Data are mean ± s.e.m; *n* = 4 biological replicates; statistical analysis performed is multiple unpaired t test with Welch’s correction. ns =not significant, ^∗^ < 0.05, ^∗∗^ < 0.01, ^∗∗∗^ < 0.001, ^∗∗∗∗^ < 0.0001. (e) Schematic of *G3BP1* minigene construct encompassing exon 2-3 and the cryptic exon. Black box depicts 150 nucleotides flanking intronic sequence included in the minigene construct. TDP-43 TG-rich binding motifs are highlighted in purple and SRRM4 TGC binding motifs in green. Stripped box in exons 2 and 3 represent minigene specific sequence. Dotted lines represent splice sites used in the case of cryptic inclusion/exclusion. (f) RT-PCR assay showing the effect of FLAG-SRRM4 expression upon doxycycline treatment (3µg/ml) in U2OS cells on cryptic exon inclusion in *G3BP1*. (g) RT-PCR assay showing the effect of TDP-43 overexpression in the presence or absence of FLAG-SRRM4 expression upon doxycycline treatment (+, 1µg/ml) or (++, 3µg/ml) in U2OS cells on *G3BP1* cryptic exon inclusion levels. (h) The association of *SRRM4* with *CRYPTIC G3BP1* RNA levels in counts per million (CPM) within the central nervous system (CNS) of ALS-TDP(+) patient samples is shown using a Spearman correlation test. ns =not significant, ^∗^ < 0.05, ^∗∗^ < 0.01, ^∗∗∗^ < 0.001, ^∗∗∗∗^ < 0.0001.

To further evaluate the regulation of *G3BP1* by SRRM4, we generated a minigene containing *G3BP1* exons 2 and 3 as well as the cryptic exon, with each exon surrounded by ∼150 nucleotides of flanking intronic sequence, including TDP-43 and SRRM4 binding sites (**Figure 6e**). This minigene was transiently transfected into the dox-inducible FLAG-SRRM4 U2OS cell line. In agreement with our analysis of the splicing of endogenous *G3BP1*, the cryptic exon in the *G3BP1* minigene was included only in the presence of SRRM4 expression, as assessed by RT-PCR using primers specific for the minigene (**Figure 6f**). Although SRRM4 expression in the non-neuronal cell lines assessed was sufficient for cryptic exon 2a inclusion in *G3BP1*, this occurs in the context of TDP-43 loss-of-function in neurons **(Figure 1, 2**), prompting us to assess the interplay between TDP-43 and SRRM4. We thus assessed the effect of TDP-43 levels on *G3BP1* cryptic exon inclusion in the context of our minigene assay. We observed that FLAG-TDP-43 transient overexpression reduced the level of cryptic *G3BP1* exon inclusion when U2OS cells were induced to express low (1µg/ml dox) or higher (3µg/ml dox) levels of SRRM4 (**Figure 6g**). These results suggest that TDP-43 and SRRM4 competitively regulate *G3BP1* splicing, acting to repress and enhance cryptic exon inclusion, respectively.

Finally, we then examined whether there is a relationship between the levels of *SRRM4* and *CRYPTIC G3BP1* in the context of ALS patients using the NYGC RNA-seq dataset. We observed a significant correlation between *SRRM4* and *CRYPTIC G3BP1* mRNA levels within the central nervous system (CNS) of ALS patients with TDP-43 pathology (**Figure 6h**). These results suggest that the relative levels of SRRM4 modulate *CRYPTIC G3BP1* burden.

Taken together, these findings support a model where TDP-43 nuclear depletion in ALS patients unmasks the SRRM4 binding site in intron 2 of *G3BP1*, driving cryptic exon2a inclusion.

Importantly, SRRM4 is a neuronal enriched splicing factor highlighting the enhanced vulnerability of neurons implicated in ALS and establishes a mechanism for the cell type variability observed in cryptic splicing signatures upon TDP-43 loss-of-function.

## DISCUSSION

In the present study, we utilize human iPSC-derived neurons and a large multi-omics dataset of ALS/FTD patients to demonstrate that TDP-43 pathology is associated with the inclusion of an in-frame cryptic exon in human *G3BP1.* We show that the cryptic peptide of 10 amino acids inserts within the highly conserved NTF2L domain of G3BP1 and disrupts protein interaction networks required for stress granule assembly. Indeed, we show that CRYPTIC G3BP1 is incapable of nucleating stress granules during oxidative stress and delays stress granule formation during heat stress. We demonstrate that CRYPTIC G3BP1 can associate with WT G3BP1 and act as dominant negative to prevent stress granule formation. We further identify a mechanism for neuronal vulnerability in ALS/FTD by elucidating how decreased levels of nuclear TDP-43 unmasks an SRRM4 binding site within *G3BP1* intron 2, allowing the inclusion of the cryptic exon2a in *G3BP1*. Our findings are consistent with a model where TDP-43 pathology involves a gain-of-function of the neuronal splicing regulator SRRM4 to include cryptic exons in transcripts and, specifically, the cryptic exon induced in *G3BP1* impairs neuroprotective stress granule assembly.

Previously we reported that TDP-43 regulates *G3BP1* mRNA stability by binding a GU-rich element within its 3’ UTR [30]. We now define a mode for *G3BP1* pre-mRNA splicing regulation by TDP-43. We delineate this by showing that a modest decrease in TDP-43 in i^3^-cortical neurons, using a low dose of TDP-43-targeting ASO, does not affect global *G3BP1* mRNA levels but allows for the inclusion of cryptic exon2a in *G3BP1*. Using a higher dose of TDP-43 ASO, we observe both modes of regulation: the inclusion of the cryptic exon in *G3BP1* and decreased *G3BP1* mRNA stability. Interestingly, cryptic exon inclusion in *STMN2* [15–17] and *UNC13A* [13, 14] upon TDP-43 depletion generates a truncated mRNA with a premature polyadenylation site and a premature termination codon, respectively, generating unstable mRNAs degraded by RNA surveillance mechanisms [13–17]. In contrast, the cryptic exon inclusion in *G3BP1* occurs in-frame and does not trigger NMD, akin to *CRYPTIC HDGFL2* [56]. However, although a significant loss in TDP-43 levels results in reduced levels of *G3BP1* mRNA and protein, no change has been reported for HDGFL2 protein levels [56]. To our knowledge, *G3BP1* is the only target of TDP-43 to be post-transcriptionally regulated by two independent mechanisms, potentially underscoring its physiological relevance in TDP-43 proteinopathies.

Cryptic splicing has been recently described as an early event in ALS, coinciding with TDP-43 nuclear redistribution and preceding clinical symptom onset [12, 56]. Conversely, more extensive cytoplasmic accumulation/aggregation of TDP-43 occurs later in ALS and correlates with clinical manifestation [12]. Given that TDP-43-dependent splicing and mRNA stability mechanisms respond differently to the degree of TDP-43 loss-of-function, our findings suggest that *G3BP1* exhibits stage-specific dysregulation in ALS. Specifically, at an early stage of the disease low levels of CRYPTIC G3BP1 are sufficient to function as a dominant negative, whereas at later stages a global reduction in *G3BP1* mRNA and protein levels will ensue, both converging on impaired stress granule formation (**Figure 7**). Several studies underscore a neuroprotective role for stress granules [22, 24–26, 57], herein our results favor this model whereby compromised stress granule function in neurons in patients with TDP-43 proteinopathies increases their susceptibility to environmental stressors implicated in disease progression.

**Figure 7:**
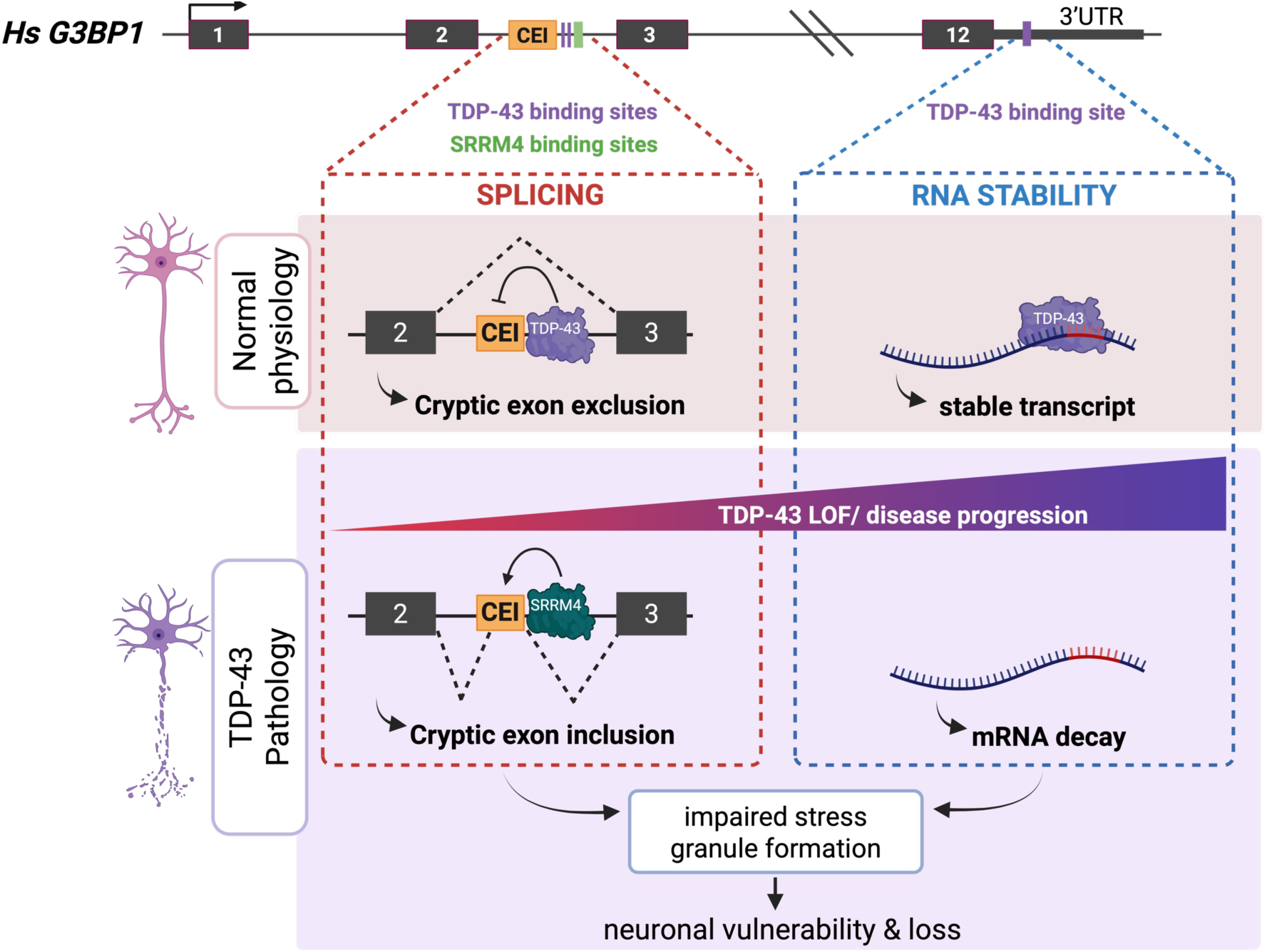
Mechanistic model for TDP-43 pathology unmasking SRRM4-mediated cryptic exon in *G3BP1*.

G3BP1 and G3BP2 function as central nodes within the stress granule protein interaction network owing to their high valency [41, 42]. The majority of G3BP1’s valency arises from protein interactions mediated by its NTF2L domain [41, 42]. Our proteomics results show that the cryptic peptide insertion in the NTF2L domain of G3BP1 results in reduced protein interactions with numerous core stress granule components, including CAPRIN1. Notably, these interactions were lost under basal, non-stressed conditions. Submicroscopic, pre-existing stress granule interaction networks are present in unstressed cells, referred to as “pre-stress seeds”, that are poised to rapidly phase separate into visible stress granules upon exposure to stress [47, 51, 58]. Thus, our proteomics data suggest that CRYPTIC G3BP1 fails to efficiently form pre-stress seeds under basal conditions. Indeed, compared to WT G3BP1, CRYPTIC G3BP1 complementation in U2OS^ΔΔG3BP1/2^ cells failed to rescue stress granule nucleation upon oxidative stress and delayed stress granule formation under heat stress. In addition, we show that CRYPTIC G3BP1 functions as a dominant negative under oxidative stress. We posit that the association of endogenous WT G3BP1 with CRYPTIC G3BP1 reduces their combined valency. Taken together, our results demonstrate that the cryptic peptide in G3BP1 perturbs its connectivity within the stress granule network, altering its capacity to form functional complexes required for stress granule assembly.

Cryptic splicing due to TDP-43 loss-of-function varies substantially across cell types but why this occurs is not known [10, 20]. In this study, we show that the cryptic splicing of *G3BP1* occurs in neurons due to its regulation by the neural-enriched splicing factor SRRM4 (or nsR100). Specifically, we show that TDP-43 and SRRM4 compete for splice site selection in intron 2 of *G3BP1*, and that *SRRM4* levels correlate with *CRYPTIC G3BP1* in ALS patients with TDP-43 pathology. These results suggest that TDP-43 pathology unmasks an SRRM4 binding site within *G3BP1* allowing for cryptic exon inclusion. Interestingly, SRRM4 further enhanced cryptic exon inclusion in *STMN2*, but not in *UNC13A*. It is possible that the presence of an SRRM4 binding motif embedded within the TDP-43 binding sites in *G3BP1* and *STMN2*, but not in *UNC13A*, supports this competitive mechanism in the former but not in the latter, although this remains to be tested **(Figure S6g, l).** SRRM4 has been shown to regulate the inclusion of microexons, typically 3-27 nucleotide in size, and this is dysregulated in autism spectrum disorder (ASD) where brains of patients show reduced expression of *SRRM4* [59]. Given that we exemplify TDP-43 loss-of-function as a gain-of-function of SRRM4 in ALS/FTD, we posit that there may be a network of transcripts/microexons dually regulated by TDP-43 and SRRM4. Recently, the small molecule T5985871 has been identified as a microexon inhibitor, acting to decrease *SRRM4* mRNA expression [60]. Whether such inhibitors can mitigate cryptic exon inclusions in TDP-43 proteinopathies warrants further investigation.

TDP-43 pathology has been reported in other cell types, albeit less frequently, such as oligodendrocytes and astrocytes [61, 62], which lack SRRM4. How TDP-43 interfaces with other splicing factors in these cell types to regulate cryptic splicing and how this renders them differentially susceptible to TDP-43 pathology remains to be established. Adding to the complexity of this regulatory framework, TDP-43 is re-localized within the nucleus under different stresses [63, 64], and TDP-43 cytoplasmic mislocalization can co-sequester certain splicing regulators [65]. It will be important to elucidate how this changes the ability of TDP-43 to interface with competing splicing regulators and their common targets. Our work herein illuminates the intricate complexity of TDP-43–mediated cryptic splicing repression with important implications for therapeutic development for TDP-43 proteinopathies.

## MATERIALS AND METHODS

### Cell culture and transfection

HeLa, HEK293FT, U2OS parental and U2OS^ΔΔG3BP1/2^ cells were grown in Dulbecco’s modified Eagle’s medium (DMEM, Wisent, 319-005-CL) supplemented with 10% fetal bovine serum (Bio-Techne, S11550H), 50 U/ml penicillin, and 50 mg/ml streptomycin at 37 °C, 5% CO_2_. Doxycycline inducible cell lines were grown in Dulbecco’s modified Eagle’s medium (DMEM) supplemented with 10% TET-FREE fetal bovine serum (TAKARA, 631101). All cells were routinely tested negative for mycoplasma.

For transient expression, plasmids were transfected using Lipofectamine LTX (Thermofisher 15338-100) and harvested after 24 hours. siRNA against human TDP-43 (L-012394-00-0005) and non-targeting negative control (D-001810-10-20) were purchased from Dharmacon. Cells were transfected with siRNA (120 nM final concentration) using RNAimax (Thermofisher, 13778150) for 5 hours and harvested after 72 hours. Lentivirus was produced in HEK293FT cells by transfecting pLD-puro or pLEX-constructs with psPAX2 and pMD2.G plasmids. For U2OS cells, filtered lentivirus was added for two days and selected with puromycin (1µg/ml) for a week. For i^3^-cortical neurons, filtered lentivirus was concentrated using Lenti-X concentrator (Takara, 631231) and then diluted in cortical neuron media (see below) and added to neurons at day 15 for four days.

### Culturing human iPSCs and integration of NGN2

iPSCs were cultured as previously described [66]. In brief, KOLF2.1 [67] cells were cultured on Synthemax® II-SC Substrate (Corning, 3535) -coated plates and maintained in complete StemFlex media (Gibco, A3349401) by daily media changes. Colonies were washed with 1X-D-PBS and split with ReleSR (StemCell Technologies, 100-0484) onto fresh, coated plates every 3-4 days with complete media changes daily.

For tet-inducible NGN2 (neurogenin-2) integration, WT KOLF2.1J cells were washed with 1X-D-PBS and split with accutase (Gibco, A1110501). A single-cell suspension was plated on Synthemax® II-SC substrate coated plates with StemFlex media supplemented with 10μM Rho-associated protein kinase (ROCK) inhibitor Y-27632 (RI) (Gibco, A2644501). The next day, Lipofectamine Stem (Invitrogen, STEM00001) and opti-MEM reduced serum media was used for transfection of piggybac-tetO-hNGN2 plasmid along with K13 EFa1-Transposase for integration into the genome. On the following day, the media was changed to complete StemFlex media with 0.5 ug/ml puromycin (Gibco, A1113803) for selection of transfected cells. Positively transfected cells have BFP fluorescence, and with a gradual increase in puromycin (up to 12 ug/ml) non-transfected cells were eliminated in 72 hours. This was verified by microscopic evaluation of live cells to ensure the plate did not contain negative (non-BFP fluorescent) colonies. This allowed survival and proliferation of only the hNGN2 transfected cells in a polyclonal manner. Puromycin selection was continued for a total of 14 days. Post 14 days, cells were split with accutase, and frozen down for stocks in knockout serum replacement media (Gibco, 10828028). Colonies were split into fresh coated plates with ReleSR when they reached 70-80% confluence, during the selection period.

### Differentiation of iPSCs to cortical neurons

Differentiation of KOLF2.1J iPSCs into cortical neurons was performed as previously described [67]. Briefly, KOLF2.1J iPSC harboring tet-inducible-NGN2 were washed with 1X PBS and dissociated with accutase. Cells were seeded on cultrex-coated plates (R&D Systems, 3434-001-02) (Day 0) with induction media [Knockout DMEM/F12 (Gibco, 11330032), 1X N2 Supplement (Gibco, 17502048), 1X Non-Essential Amino Acids (Gibco-11140050), 1X Glutamax (Gibco 35050061] along with ROCK inhibitor Y-27632 and 2 μg/mL doxycycline (Sigma, D9891-1G) for neural progenitor cell (NPC) generation. Cultrex was diluted in DMEM/F12 (Gibco, 11320033). Complete media change was provided the next day and fresh induction media added with doxycycline. The following day, 48 hrs from plating iPSC for NPC induction, complete media change was performed with doxycycline for NGN2 induction. The following day (72 hours post seeding), the NPCs (with neurites clearly visible), were washed with 1X-D-PBS, and dissociated with accutase treatment followed by freezing down in cryovials for future plating for i^3^ cortical neurons.

Plates were coated with poly-ornithine (Sigma, P3655) diluted in borate buffer, overnight at 37°C. Next day, the plates were washed with sterile water four times and dried inside the hood. Half-volume of cortical neuron media [Brainphys (STEMCELL Technologies, 05790), 1X B27 Supplement (Gibco, A1413701), 10 ng/mL BDNF (PeproTech, 450-02), 10 ng/mL NT3 (PeproTech, 450-03), 1 μg/mL Laminin (Gibco, 23017015)] was added to the plates and kept for 1 hour before NPCs were seeded. Frozen NPCs were thawed, spun down in cortical neuron media and resuspended in fresh complete cortical neuron media and added dropwise to the prepared plates, swirled gently and incubated at 37°C . Media supplementation was done the following day. Half-media changes were given every 2-3 days.

### ASO treatment in i^3^ cortical neurons

TDP-43 ASO and non-targeting (NTC) antisense oligonucleotides (ASOs) were generously provided by Ionis Pharmaceuticals. On day 30 of differentiation, ASOs were added to culture media at a final concentration of 5 or 10 μM. Half media change was provided 3 days later. On day 37 of differentiation, a second dose of ASO was added to culture media at a final concentration of 5 or 10 μM. Half media change was provided 3 days later, and cells were harvested at day 44 of differentiation.

### iPSC derived-direct induced motor neuron (diMNs) culture

Sporadic ALS (sALS), C9orf72, and non-neurological control patient iPSC lines were acquired from the Answer ALS repository at Cedars Sinai and differentiated into mixed spinal neuron cultures as recently described [68]. Metadata for all iPSC lines used is reported in [68]. All iPSC lines used in this study were differentiated in a single batch. All cell lines were maintained at 37°C with 5% CO_2_ and routinely tested negative for mycoplasma.

Briefly, iPSCs were maintained in 6 well plates coated with growth factor reduced Matrigel (Corning, 354230) and mTeSR Plus Media (StemCell Technologies, 100-0276). iPSCs were cultured and expanded for 3 weeks and on day 0, iPSCs were dissociated with StemCell Technologies, 05872) and plated in 6 well plates coated with growth factor reduced Matrigel at a density of 5×10^5^ cells/well in stage 1 media (47.5% IMDM (Gibco, 12440053), 47.5% F12 (Gibco, 11765062), 1% NEAA (Gibco, 11140-050), 1% Pen/Strep, 2% B27 (Invitrogen, 17504044), 1% N2 (Life Technologies, 17502048), 0.2 μM LDN193189 (Stemgent, 04-0074-02), 10 μM SB431542 (StemCell Technologies, 72234), and 3 μM CHIR99021 (Sigma Aldrich, SML1046) supplemented with 20 μM ROCKi (Millipore, Y-27632). The next day, media was exchanged with stage 1 media without ROCKi. Media was subsequently exchanged on days 3 and 5 of differentiation. On day 6 of differentiation, cells were dissociated with accutase (Gibco, A11105) and plated in T25 flasks coated with growth factor reduced Matrigel at a density of 2×10^6^ cells/flask in stage 2 media (47.5% IMDM, 47.5% F12, 1% NEAA, 1% Pen/Strep, 2% B27, 1% N2, 0.2 μM LDN193189, 10 μM SB431542, 3 μM CHIR99021, 0.1 μM all-trans RA (Sigma

Aldrich, R2625), and 1 μM SAG (Cayman Chemicals, 11914) supplemented with 20 μM ROCKi. The next day, media was replaced with stage 2 media without ROCKi. Media was subsequently exchanged on days 9 and 11 of differentiation. On day 12 of differentiation, cells were dissociated with Trypsin and plated in 6 well plates coated with growth factor reduced Matrigel at a density of 5×10^6^ cells/well in stage 3 media (47.5% IMDM, 47.5% F12, 1% NEAA, 1% Pen/Strep, 2% B27, 1% N2, 0.1 μM Compound E (Millipore, 565790), 2.5 μM DAPT (Sigma Aldrich, D5942), 0.1 μM db-cAMP (Millipore, 28745), 0.5 μM all-trans RA, 0.1 μM SAG, 200 ng/mL Ascorbic Acid (Sigma Aldrich, A4544), 10 ng/mL BDNF (PeproTech, 450-02), and 10 ng/mL GDNF (PeproTech, 450-10) supplemented with 20 μM ROCKi. The next day, media was replaced with stage 3 media without ROCKi. Media was exchanged on day 16 of differentiation. On day 18 of differentiation, cells were dissociated with accutase and plated in 6 well plates coated with growth factor reduced Matrigel at a density of 5×10^6^ cells/well in stage 3 media supplemented with 20 μM ROCKi. The next day, media was replaced with stage 3 media supplemented with 20 μM AraC. The next day, media was replaced with stage 3 media without AraC. Media was subsequently exchanged every 3-4 days until day 32 of differentiation. Beginning on day 32 of differentiation, a half volume of stage 3 media was exchanged every 3-4 days until day 53 of differentiation. On day 53 of differentiation, cells were dissociated with accutase and plated in glass bottom 24 well plates coated with growth factor reduced Matrigel at a density of 3.5 ×10^5^ cells per well in stage 3 media supplemented with 20 μM ROCKi. The next day, media was replaced with stage 3 media without ROCKi. A half volume of stage 3 media was subsequently exchanged on day 57 of differentiation. Cells were harvested at day 60 of differentiation.

### RIPA lysis of i^3^ cortical neurons

Cell pellets were resuspended in RIPA lysis buffer supplemented with protease inhibitors (LPC) Leupeptin (Bioshop, LEU001), Pepstatin A (Bioshop, PEP605), Chymostatin (Sigma, C7268) and Halt phosphatase inhibitor (ThermoFisher, 78420) and kept on ice for ten minutes. The resuspended pellets were vortexed and incubated at RT for another 10 mins followed by centrifugation at 13,000 rpm for 15 mins at 4°C. The supernatant was collected and protein concentration estimated by BCA (ThermoFisher, PI23227).

### Co-immunoprecipitation

For co-immunoprecipitation (Co-IP) experiments, U2OS^ΔΔG3BP1/2^ cells were transiently transfected with HA-WT or CRYPTIC G3BP1 constructs and/or GFP-WT G3BP1 using PEI reagent (Polysciences, 23966-100). Cells were harvested 48 hours later and lysed in lysis buffer (50 mM HEPES pH 7.5, 100 mM KCl, 2 mM EDTA, 10% glycerol, 1 mM DTT, and 0.1% NP-40) supplemented with protease inhibitors (LPC), and subjected to three freeze/thaw cycles. Lysates were clarified by centrifugation at 20,000 g for 20 min at 4°C. Lysates were pre-cleared with 20 ul of Protein A dynabeads (Thermofisher, 10002D) for 30 min at 4°C with end-over-end rotation. In the meantime, 30 ul of Protein A dynabeads per sample were incubated with 3 µg of antibody indicated (G3BP1 Proteintech, 13057-2-AP, or HA abcam, ab9110) in PBS for 1 hr at room temperature with end-over-end rotation to conjugate, and then washed thrice in lysis buffer. 1.5 mg of pre-cleared lysate (quantified using Bradford assay (Biorad, 5000006) per condition was then incubated with conjugated Dynabeads in the presence of benzonase nuclease (1ul/mg of lysate, Millipore, 70746-3) for 3 hours at 4°C with end-over-end rotation. The beads were then washed five times with 1 ml of cold lysis buffer and eluted by boiling with 40 ul of 2x Laemmli sample buffer. Proteins were separated by SDS–PAGE followed by Western blotting.

### RNA extraction and RT-qPCR

RNA was extracted using RNAeasy^®^ Minikit (Qiagen) and treated with turbo DNase I (Ambion, AM1907) according to the manufacturer’s instructions. Equal amounts of RNA (500-1000ng) were reverse transcribed using the QuantiTect Reverse Transcription kit (Qiagen, 205313). PrimeTime Standard qPCR assays (Table S3) with IDT gene expression mix (IDT, 1055772) were used for quantitative PCR (qPCR) assays using the QuantStudio 7 Flex Real-Time PCR System (Life Technologies). Data was analyzed using the 2^−ΔΔCT^ method and normalized to the mean of two housekeeping genes (*GAPDH* and *18S*).

### Live imaging during oxidative stress

We seeded 1.2-1.5 ×10^4^ GFP-WT or CRYPTIC G3BP1-expressing U2OS^ΔΔG3BP1/2^ cells/well 24 hours before the experiment in a 96-well plate (Brooks, MGB096-1-2-LG-L). Then, we replaced the media with 250uM NaAsO_2_ (sodium arsenite) (Sigma, S7400) -containing medium and immediately examined them under the microscope to monitor stress granule formation. We utilized a PCO-Edge sCMOS camera controlled by VisView installed on a VisiScope Confocal Cell Explorer system (Yokogawa spinning disk scanning unit; CSU-W1) and an inverted Olympus microscope (60× oil objective; excitation wavelength: GFP: 488 nm). We analyzed stress granule and cell areas using surface features in Imaris software 9.5.1. The stress granule area per field of view (FOV) for each time point was normalized to the cell area of the same FOV as calculated from the first time point (“0 min”). Four FOVs per well were imaged. Each condition was done in four replicate wells in three independent biological replicates.

### Live imaging during heat stress

1.5 ×10^5^ cells/well were seeded into optical 24-well plates (Cellvis, P24-1.5H-N). Live imaging was performed in a Zeiss CD7 microscope. Stress was induced by raising the temperature of the microscope incubator to 43°C. Stress granules were marked at each time point of the imaging run using Imaris image analysis software. Stress granule number/cell number was computed by dividing the number of stress granules at each time point by the number of cells in the field.

### Immunofluorescence

U2OS parental cells were seeded on cover slips in a 6 well plate and transfected with 200ng HA-tagged CRYPTIC G3BP1 the following day. 24 hours post-transfection, the media was changed, and cells were treated with 0.5 mM of sodium arsenite (Sigma Aldrich) for 30 mins at 37°C. Cells were fixed with 4% PFA in PBS and washed once with 1X PBS. Cells were then permeabilized using 0.1% Triton^TM^ X-100/PBS for 15 minutes. The cells were washed thrice with 1X PBS and then blocked with Superblock (Scytek Laboratories, AAA500) at RT for 1 hour. Cells were then incubated with specific primary antibodies diluted in Superblock at RT for 1 hour. Cover slips were washed thrice with 1X PBS and incubated with secondary antibody diluted in Superblock at RT for 1 hour. Cells were then washed thrice with 1x PBS and incubated for 5 minutes with Hoechst / DAPI for nuclear staining. Coverslips were washed thrice with 1X PBS and mounted on glass slides using Prolong anti-fade (Invitrogen, P36930). Images were acquired on Leica SP5 confocal microscope using the 63x – oil objective lens. Acquired images were analyzed using LAS AF software (Leica).

### APEX proximity labeling

G3BP1-WT APEX and NES-APEX expressing U2OS^ΔΔG3BP1/2^ cell lines were induced with 25ng/ml tetracycline (Sigma, T7660) in order to match the expression level and biotiniylation activity level of the G3BP1-CRYPTIC APEX line. Cells were seeded into T80 flasks with tetracycline 24 hours before APEX activation. Biotin-Tyramide (BP) (Iris Biotech, 41994-02-9) was added to a final concentration of 500uM and incubated for 1 hour at 37°C. Then H_2_O_2_ (JT Baker, 2186-01) was added to a final concentration of 1mM for exactly 60 seconds to activate APEX activity in a temporally controlled manner. APEX activity was then quenched using quencher solution (10mM sodium ascorbate (Alfa Aesar, A17759), 5mM Trolox (Sigma, 238813) and 10mM sodium azide (Sigma, S2002) in PBS). Cells were harvested by scraping in quencher solution, centrifuged and frozen in liquid nitrogen. Cells were lysed in RIPA buffer (50mM Tris-HCl pH8, 150mM NaCl, 1% NP-40, 0.5% sodium deoxycholate, 0.1% SDS) and centrifuged at 15000g at 4°C for 10 min. 0.5mg of lysate for each sample was incubated with 100ul streptavidin magnetic beads (Pierce Streptavidin Magnetic Beads, ThermoFisher, 88817) overnight at 4°C with rotation. Beads were then washed twice with RIPA buffer, once with 1M KCL, once with 0.1M Na_2_CO_3_ and once with 2M urea/10mM Tris-HCl pH8.

### Mass spectrometry

The samples were subjected to on-bead tryptic digestion. The resulting peptides were analyzed using nanoflow liquid chromatography (nanoAcquity) coupled to high resolution, high mass accuracy mass spectrometry (Q-Exactive HFX). Each sample was analyzed on the instrument separately in a random order in discovery mode. The raw data were processed with MaxQuant v1.6.6.0. The data were searched with the Andromeda search engine against the human proteome database as downloaded from Uniprot, appended with common lab protein contaminants. Search parameters included the following modifications: Fixed modification-cysteine carbamidomethylation. Variable modifications-methionine oxidation, asparagine and glutamine deamidation, and protein N-terminal acetylation.

### Proteomics analysis

MaxQuant proteinGroups output was analyzed using Perseus. Proteins detected as contaminants were first removed. Intensity values were log_2_ transformed and missing values were imputed for proteins with at least four valid values using an artificial normal distribution with a downshift of 1.8 standard deviations and a width of 0.3 of the original ratio distribution. Student’s t test with FDR <=0.05 was used to compare each sample to no BP control samples to identify proteins non-specifically bound to streptavidin beads. These were filtered out and proteins were then compared between G3BP1-APEX and NES-APEX baits. LFQ (label-free quantification) values were log_2_ transformed and missing values were imputed for proteins with at least two valid values in at least one group using an artificial normal distribution with a downshift of 1.8 standard deviations and a width of 0.3 of the original ratio distribution. Proteins were then compared between each G3BP1-APEX bait to the NES-APEX bait using Student’s t test with FDR <=0.05 to determine which proteins are significantly enriched in the vicinity of G3BP1 (either WT or CRYPTIC) compared to NES. G3BP1-APEX values were then normalized by the mean of their corresponding NES-APEX values. The NES-normalized values were then compared between the G3BP1-WT APEX bait vs G3BP1-CRYPTIC APEX bait using Student’s t test with FDR <=0.05. log2 fold change (log2FC) was computed by Mean log2 LFQ_CE−Mean log2 LFQ_WT and represented as a volcano plot using R. GO enrichment analysis was performed using ShinyGO 0.85 (http://bioinformatics.sdstate.edu/go/).

### Alphafold prediction of NTF2L CRYPTIC G3BP1 – CAPRIN1

The structure of the NTF2L domain of WT G3BP1 (a.a. 1-146) in complex with the NTF2L domain of CRYPTIC G3BP1 (a.a. 1-156) was modelled with AlphaFold 3, using a publicly available server (https://alphafoldserver.com/) [50]. The prediction produced 5 models, which converged (rmsd <0.7 Å) with an interface predicted template modelling (ipTM) score of 0.84. The top predicted structure was superposed on the crystal structure of the WT G3BP1 NTF2L dimer bound to a CAPRIN1 peptide (PDB 8TH7) using PyMOL version 3.1.4.1, which we used to create Figures 3e and 5a.

### Cloning and stable cell lines

HA-tagged WT G3BP1 was generated by subcloning from GFP-G3BP1 using restriction enzyme cloning techniques into PCl-lambda-N-HA plasmid (gift from Marc R Fabian). HA-tagged CRYPTIC G3BP1 was cloned into PCl-lambda-N-HA plasmid using g-block (IDT) and restriction enzyme cloning techniques. GFP-tagged WT and CRYPTIC G3BP1 were cloned as described previously [47] using the restriction-free cloning method. Tetracycline inducible G3BP1-APEX and NES-APEX constructs and cell lines were generated as previously described [47]. <u>pLDPuro-</u> <u>hsnSR100N</u> (FLAG-tagged SRRM4) plasmid (Addgene, 35172) was used to produce lentivirus to infect U2OS parental cells. Restriction-free cloning method using NEBuilder® HiFi DNA Assembly Cloning Kit (NEB, E5520S) was used according to the manufacturer’s instructions to clone WT or CRYPTIC G3BP1 into pLEX-PGK-EYFP backbone from pLEX-PGK-LAMP1-EYFP plasmid (Addgene, 164215). G3BP1-minigene was generated using a g-block (IDT) to comprise human exon 2, cryptic exon and exon 3, and 150bp intronic sequence flanking each exon. First/last four amino acids in exon 2/3, respectively, were wobbled to create minigene-specific sequences. Restriction enzyme cloning techniques was then used to insert into pCDNA vector.

### Minigene assay

3 ×10^5^ U2OS parental cells were seeded per well in a 6-well format. Next day cells were transfected with 2μg FLAG-TDP-43 pCDNA versus empty pCDNA control and 50ng G3BP1-minigene pCDNA. Doxycycline was added to a final concentration of 0, 1 or 3 μg/ml. Cells weren harvested after 24 hours and RNA extracted as described above. For RT-PCR assay, Q5 enzyme (NEB, M0491S) was used as per manufacturer’s instructions to amplify PCR product using primers specific for minigene and was run on a 3% agarose gels containing GelRed (Biotium, 41003).

### Bioinformatics analysis

Publicly available dataset (iPSC-cortical neuron +/- TDP-43 knockdown) was obtained from European Nucleotide Archive (ENA) under accession PRJEB42763. To generate sashimi plot we used the aligned files available via the ENA portal and set the splicing junction minimum to 5 junction reads. To generate the sashimi plot we used the aligned files (hg38) available via the ENA portal. To better visualize the data and reduce background noise, we set the splicing junction minimum to 5 junction reads. TDP-43 (TARDBP, K562) eCLIP data is available at ENCODE (doi:10.17989/ENCSR720BJU) and was visualized in Integrative Genomics Viewer (IGV) [69]. Sequence conservation of *G3BP1* cryptic exon2a was assessed using the Primates Multiz Alignment & Conservation track on UCSC genome browser [70].

### New York Genome Center ALS Consortium (NYGC) dataset

Our analysis contains transcriptomics data from 2361 tissues from 640 individuals with ALS or FTD versus non-neurological controls. To assess the impact of TDP-43 pathology on cryptic splicing, we performed an analysis comparing patients with TDP-43 pathology to TDP-43 pathology-free samples (non-TDP controls), that were either controls or patients harboring *SOD1* or *FUS* mutations. Sample processing, library preparation, and RNA-seq quality control and alignment for the NYGC data have been described in previous papers [13, 71, 72]. Splice junction reads for the cryptic splice junctions queried were extracted from aligned BAM files using RegTools [73] using a minimum of 8 bp as an anchor on each side of the junction and a maximum intron size of 500 kb. Counts per million (CPM) was calculated as the number of unique splice junctions mapped to the cryptic divided by the library depth of the sample BAM file. Spearman correlation analysis was performed on non-zero CPM values for *CRYPTIC G3BP1* and other cryptic exon containing transcripts within the central nervous system (CNS) of ALS/FTD patients with TDP-43 pathology.

Expression levels of the SRRM4 gene were obtained using the Feature counts tool. A count file for SRRM4 for each sample was generated and used for the next step of the analyses. SRRM4 expression was normalized to counts per million (CPM), calculated as raw counts divided by the total number of mapped reads per sample and scaled to one million, CPM=((srrm4-count/library size)*1000000) for available samples (1152 from ALS patients with TDP-43 pathology). Spearman correlation analysis was then performed on *SRRM4* and *CRYPTIC G3BP1* CPM levels within the central nervous system (CNS).

### Quantification and statistical analysis

The number of replicates as well as statistical tests performed and significance values are indicated in the figure legends. Data is presented as the mean +/- standard error of the mean (SEM). Multiple unpaired t tests, Mann-Whitney tests and Spearman correlations were performed using GraphPad Prism10 software (La Jolla, CA).

## Acknowledgments

We thank Michael E Ward and William C Skarnes for providing us with the KOLF2.1 cells and K13 EFa1-Transposase plasmid, as well as their valuable discussions. We thank Shawn M Lyons and Paul J Anderson for providing us with the U2OS^ΔΔG3BP1/2^ cells. This work was supported by Canadian Institutes of Health Research (CIHR) as well as ALS Canada in partnership with Brain Canada. HF was supported by ALS Canada and Fonds de recherche du Québec (FRQS). Schematic diagram was created in Biorender.

---

**Figure S1.**
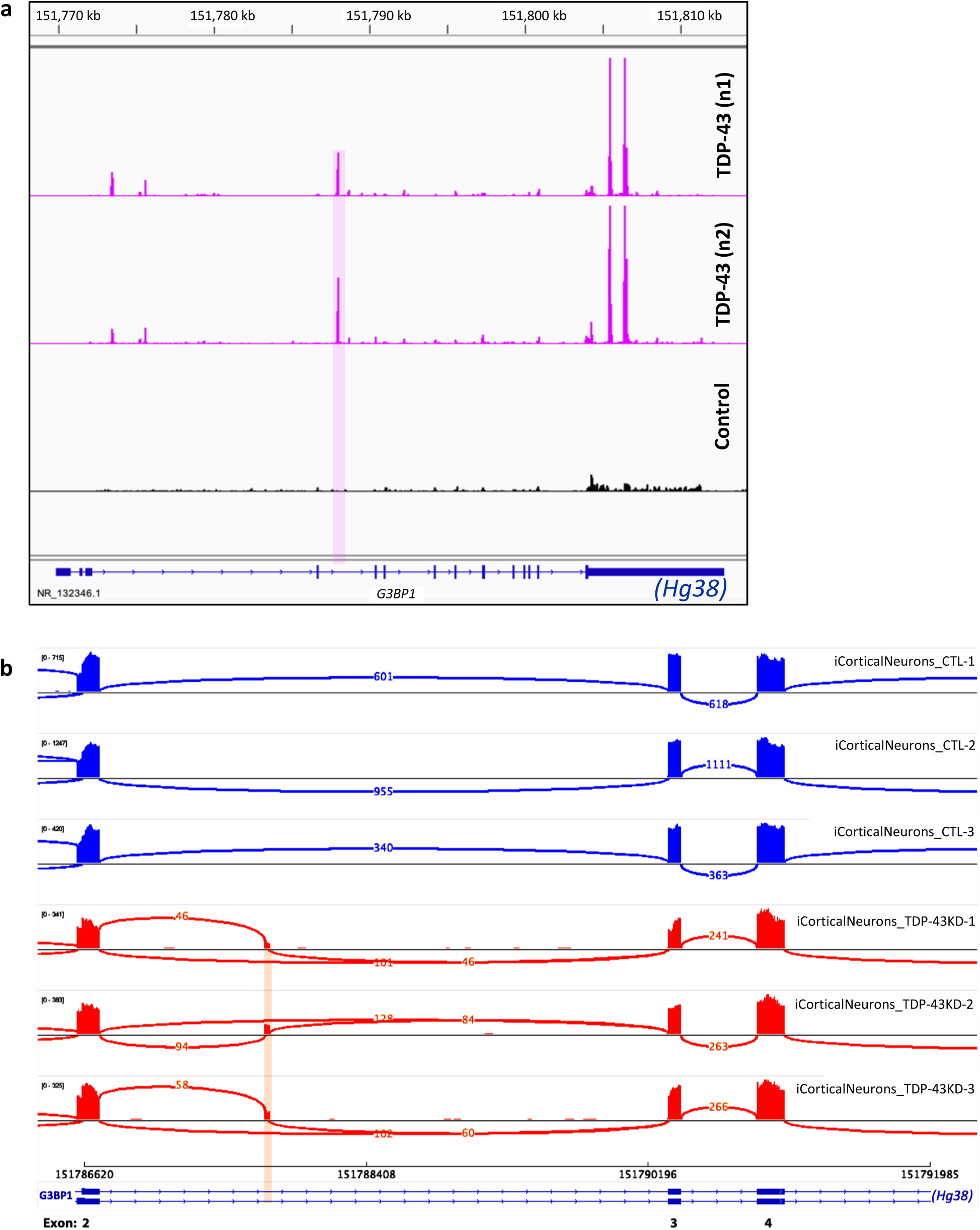
Related to Figure 1, TDP-43 depletion in cortical neurons leads to the inclusion of a cryptic exon in *G3BP1*. (a) eCLIP unique reads for TDP-43 (2 replicates) versus control showing TDP-43 binding sites localized between exon 2 and 3 (intron 2) and 3’ UTR of human *G3BP1*. (b) IGV plot for differential splicing analysis performed by MAJIQ in control (n = 3) and CRISPRi TDP-43 depleted (KD) (n = 3) iPSC-derived i^3^ cortical neurons.

**Figure S2.**
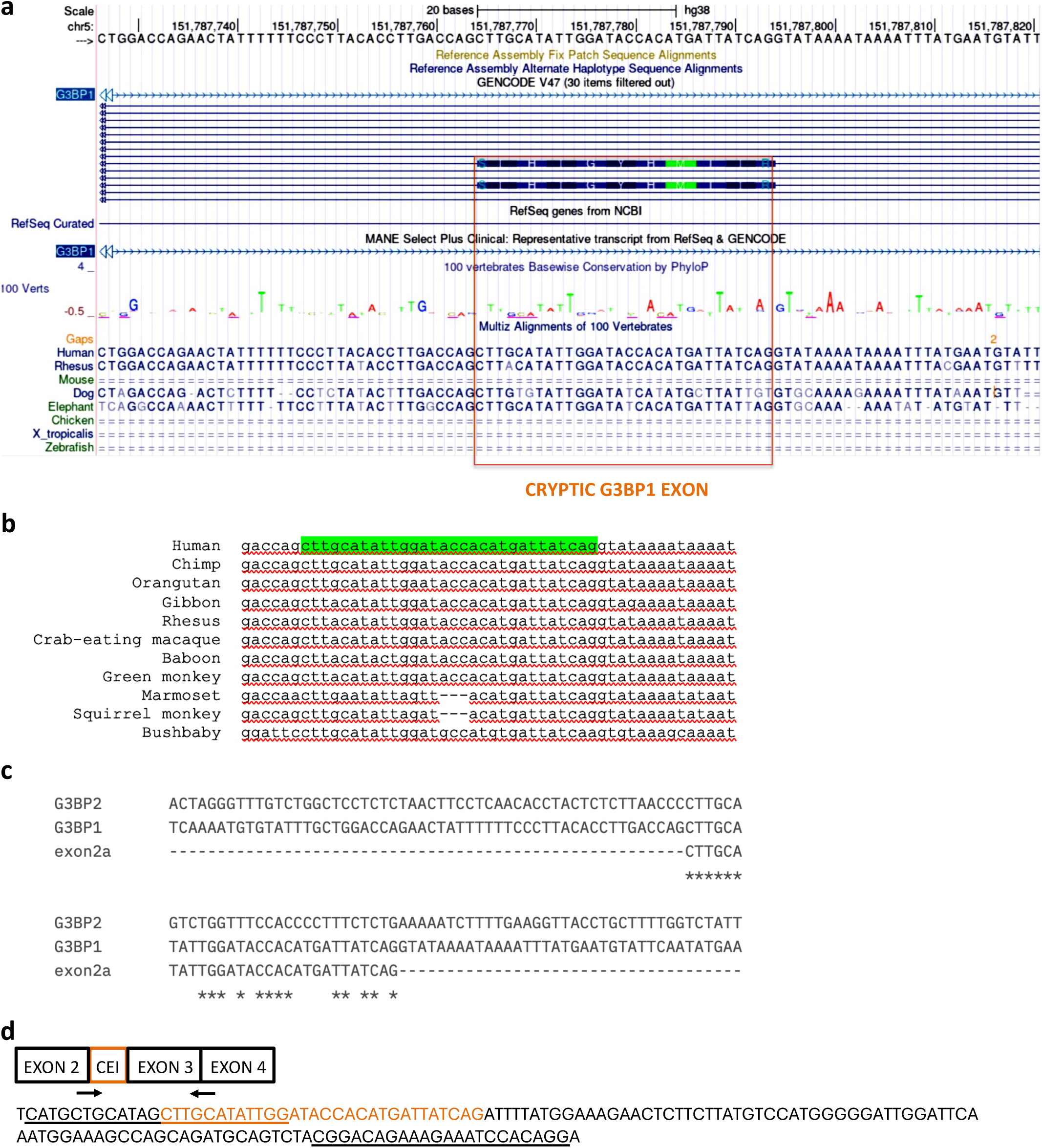
Related to Figure 1, Sequence conservation of *G3BP1* cryptic exon 2a and primer design. (a) Multiple sequence alignment using UCSD genome browser 100 vertebrates conservation track showing that *G3BP1* cryptic exon2a sequence is conserved in rhesus monkey but not in mouse or zebrafish. (b) Extended multiple sequence alignment showing *G3BP1* cryptic exon2a sequence conservation across several primates. (c) Sequence alignment of human *G3BP1*, *G3BP2*, and *G3BP1* exon2a showing lack of sequence conservation and absence of cryptic 3′/5′ splice sites in *G3BP2*. (d) Schematic of localization of RT-qPCR primer pair (arrows) used to detect *CRYPTIC G3BP1 exon2a* and (below) longread sequencing results for the qPCR amplicon for *CRYPTIC G3BP1* mRNA in i^3^ cortical neurons depleted for TDP-43.

**Figure S3.**
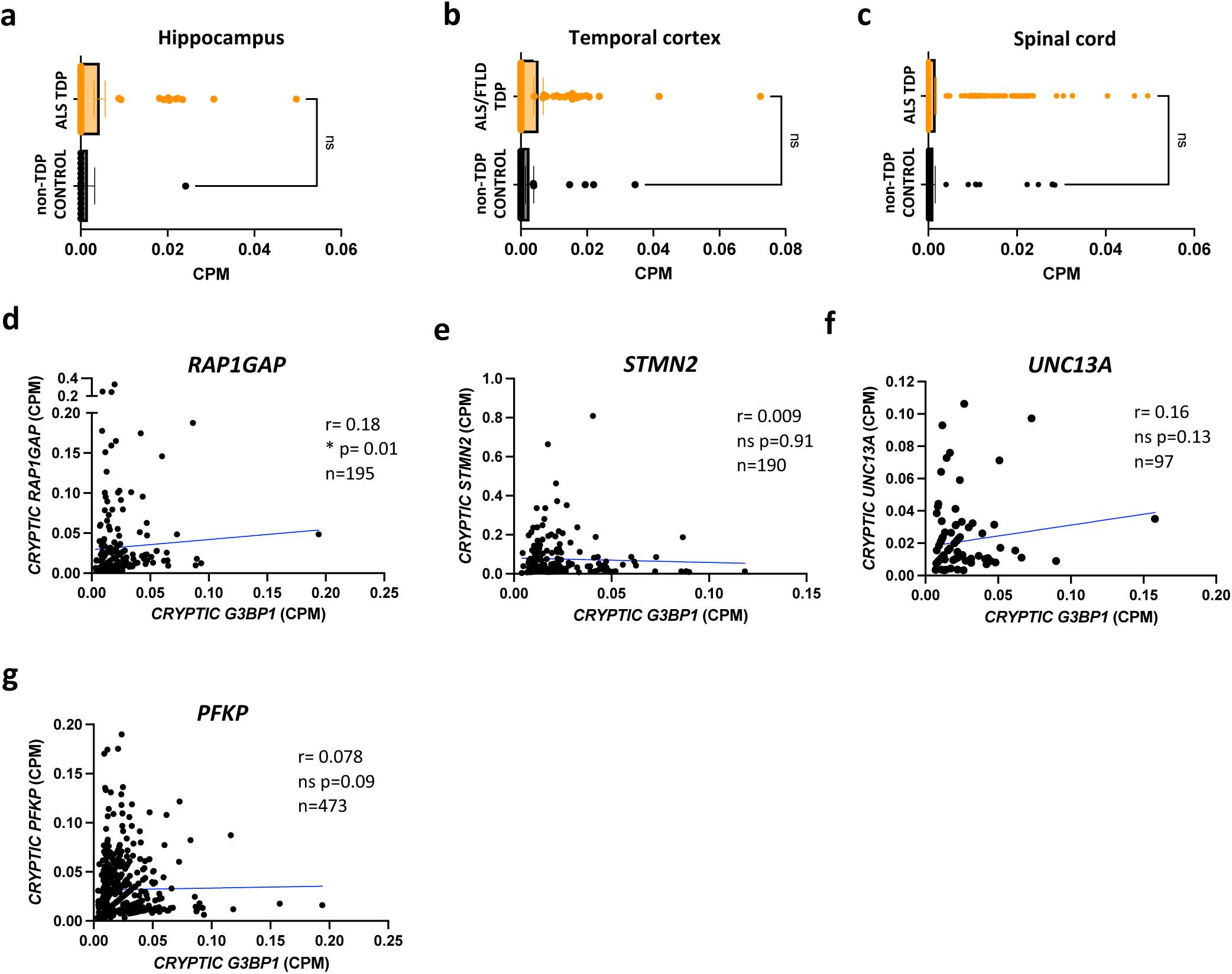
Related to Figure 2, *CRYPTIC G3BP1* is detected in ALS/FTLD patient neurons and correlates with TDP-43 pathology. (a-c) Expression of *CRYPTIC G3BP1* transcript across indicated brain regions from NYGC ALS Consortium RNA seq depicted as counts per million (CPM). Samples were stratified by TDP-43 pathology, comparing ALS/FTD patients with TDP-43 pathology (ALS/FTD TDP) to TDP-43 pathology-free samples (non-neurological controls or patients with *SOD1* or *FUS* mutations (non-TDP CONTROL)). Hippocampus (non-TDP CONTROL n=15, ALS TDP n=61), Temporal cortex (non-TDP CONTROL n=37, ALS/FTLD TDP n=68), Spinal cord (non-TDP CONTROL n=153, ALS TDP n=704). Data are mean ± s.e.m; statistical analysis performed is Mann-Whitney test. ns = not significant. (d-g) Correlation between *CRYPTIC G3BP1* levels and indicated cryptic exon-containing transcripts (in CPM) within the central nervous system (CNS) of ALS/FTD patients with TDP-43 pathology. Statistical analysis performed is Spearman correlation test. ns =not significant, ^∗^ < 0.05.

**Figure S4.**
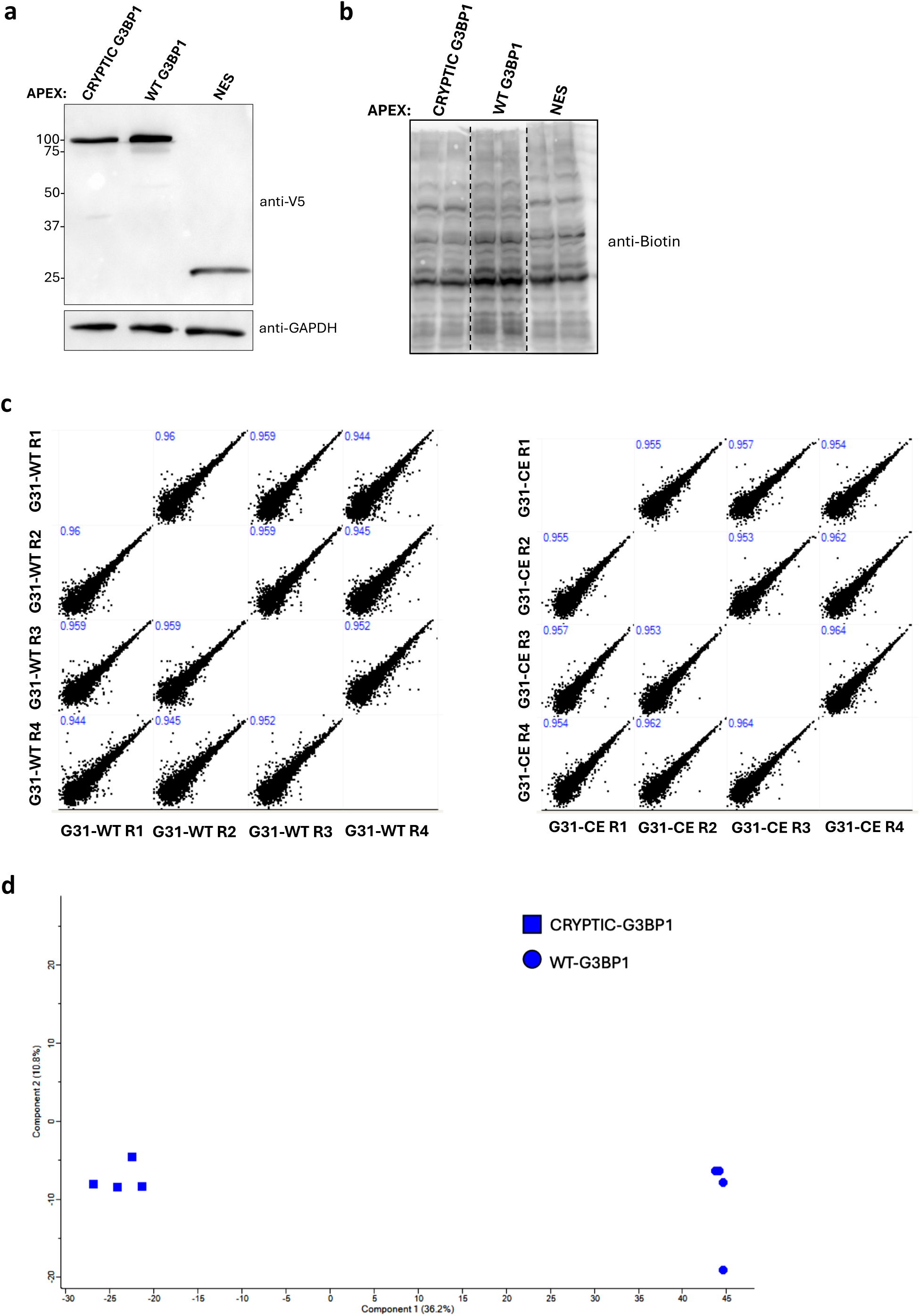
Related to Figure 3, APEX-G3BP1 experimental controls and replicates. (a) Western blot analysis showing expression levels of APEX-tagged CRYPTIC G3BP1, WT G3BP1 and NES in U2OS^ΔΔG3BP1/2^ cells. GAPDH was used as a loading control. (b) Western blot analysis probed for Biotin showing BP-tagged endogenous proteins upon activation of APEX2 in U2OS^ΔΔG3BP1/2^ cells expressing APEX-tagged CRYPTIC G3BP1, WT G3BP1 and NES. (c) Scatter plot and Pearson correlation coefficient values in between experimental replicates (APEX-WT G3BP1 on left, APEX-CRYPTIC G3BP1 on right). (d) Principal-component analysis (PCA) of APEX-WT G3BP1 and APEX-CRYPTIC G3BP1 proteomes under basal non-stressed conditions.

**Figure S5.**
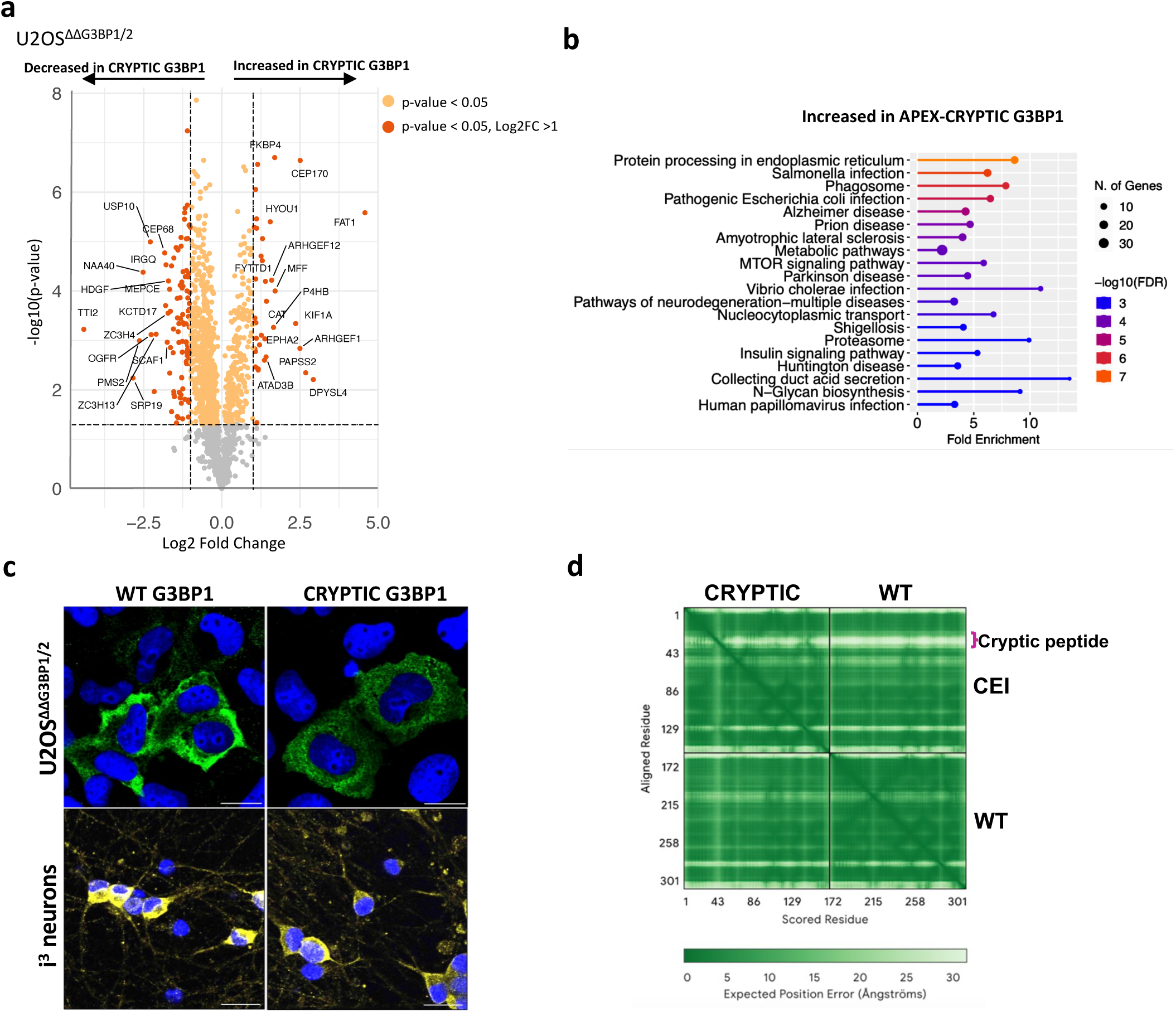
Related to Figure 3, *CRYPTIC G3BP1* adds 10 amino acids in-frame in NTF2L domain and alters its protein interactome. (a) Volcano plot of relative protein levels analyzed by mass spectrometry in APEX-CRYPTIC G3BP1 versus APEX-WT G3BP1 samples, after normalizing to NES-APEX. y axis depicts the differential expression p values (−log10 scale) with dotted horizontal line showing –log10(p=0.05) for significance. x axis depicts log2fold change (FC) with vertical dotted line showing log2FC(1). All orange dots show significance for p-value (p< 0.05), dark orange dots show genes with significant p value and a log2FC≥ +/- 1. Top 15 upregulated and downregulated genes are labeled. *n=4* biological replicates. (b) KEGG pathway analysis of significantly upregulated genes (p< 0.05) in CRYPTIC G3BP1-APEX versus WT G3BP1-APEX samples, after normalizing to NES-APEX, using ShinyGO tool. (c) Immunofluorescence of cells transiently expressing HA-WT G3BP1 or HA-CRYPTIC G3BP1 in U2OS^ΔΔG3BP1/2^ imaged using HA antibody versus iPSC-derived cortical i3Neurons stably expressing YFP-WT G3BP1 or YFP-CRYPTIC G3BP1 using YFP antibody (Scale bar, 20 μm). (d) PAE (Predicted aligned error) plot for the CRYPTIC G3BP1 NTF2L domain bound to the WT G3BP1 NTF2L. The low expected error between the two domains indicates a high confidence prediction.

**Figure S5.**
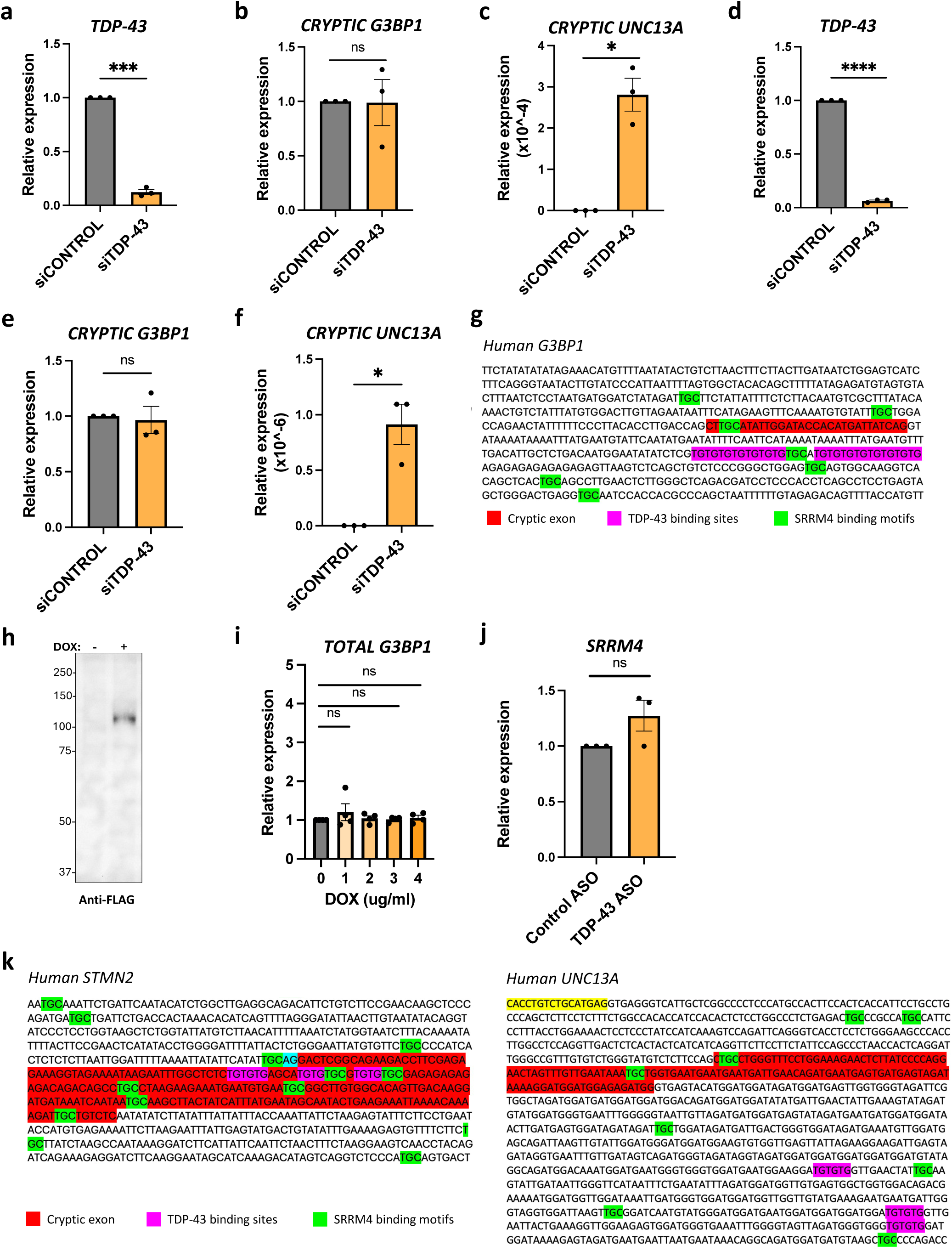
Related to Figure 6, Neuron-enriched splicing factor SRRM4/nsR100 enhances cryptic *G3BP1* exon2a inclusion and competes with TDP-43. (a-f) RT-qPCR analysis for specified gene (normalized to housekeeping genes *GAPDH* and *18S*) where (a-c) are in U2OS cells and (d-f) are in HeLa cells. (a,b,d and e) are expressed as a fold change over siControl, whereas (c) and (f) depict relative expression levels. Data are mean ± s.e.m; *n* = 3 biological replicates; statistical analysis performed is multiple unpaired t test with Welch’s correction. ns =not significant, ^∗^ < 0.05, ^∗∗^ < 0.01, ^∗∗∗^ < 0.001, ^∗∗∗∗^ < 0.0001. (g) Sequence surrounding cryptic exon in *G3BP1*. Cryptic exon is highlighted in red, TDP-43 binding sites in purple, and SRRM4 binding motifs in green. (h) Western blot analysis showing FLAG-SRRM4 expression at 3µg/ml dox concentration using FLAG antibody. (i) RT-qPCR analysis for *total G3BP1* (normalized to housekeeping genes *GAPDH* and *18S*) as a fold change over no doxycycline (dox) control. x axis shows doxycycline concentration used to induce FLAG-SRRM4 expression in stable U2OS cell line. Data are mean ± s.e.m; *n* = 4 biological replicates; statistical analysis performed is multiple unpaired t test with Welch’s correction. ns =not significant. (j) RT-qPCR analysis for *SRRM4* (normalized to housekeeping genes *GAPDH* and *18S*) as a fold change over control. 10μM of control (non-targeting) or TDP-43 ASO treatment used. Data are mean ± s.e.m; *n* = 3 biological replicates; statistical analysis performed is multiple unpaired t test with Welch’s correction. ns =not significant. (k) Sequence surrounding cryptic exon in *STMN2 (left)* and *UNC13A* (right). Cryptic exon is highlighted in red, TDP-43 binding sites in purple, and SRRM4 binding motifs in green.

